# A highly active bacterial actin actuates the polymerization of another isoform essential for swimming motility of *Spiroplasma*

**DOI:** 10.1101/2024.09.04.611326

**Authors:** Daichi Takahashi, Hana Kiyama, Hideaki T. Matsubayashi, Ikuko Fujiwara, Makoto Miyata

**Author notes:** Research Institute for Interdisciplinary Science, Okayama University, Okayama, Japan (Daichi Takahashi).

## Abstract

*Spiroplasma* is a wall-less helical bacterium that is characterized by a unique swimming motility involving five isoforms of the bacterial actin MreBs (SMreB1–5). The functions of SMreBs are unique in the MreB family proteins, as their counterparts in walled-bacteria are involved in maintaining the cell shape by scaffolding the cell-wall synthesis complex through their static properties. *In vitro* analyses of individual SMreBs provide clues to understand the detailed molecular mechanism of *Spiroplasma* swimming. However, the purification difficulties have hampered *in vitro* analyses of one of the SMreBs, SMreB1, which drives the swimming. Here, we isolated soluble SMreB1 of *Spiroplasma eriocheiris* (SpeMreB1) and evaluated its activity. SpeMreB1 was expressed as a fusion with a solubilization-tag, ProteinS (PrS), which allowed us to purify it in the soluble fraction. SpeMreB1 exhibited the highest phosphate release rate and the fold changes of critical concentrations for polymerization across the nucleotide states among the MreB family proteins. SpeMreB1 interacted with polymerized SpeMreB5, another SMreB essential for *Spiroplasma* swimming. In the AMPPNP- or ADP-bound state, SpeMreB1 decreased the amount of SpeMreB5 filaments, possibly reflecting their disassembly. Regardless of the nucleotide state, SpeMreB1 bound to negatively charged lipids. These results suggest that SpeMreB1 utilizes its highest activity to manage SpeMreB5 filaments underneath the cell membrane to drive *Spiroplasma* swimming.

**Importance:** In most bacterial species, MreB is involved in cell-shape maintenance by localizing the bacterial cell-wall synthesis complex with its static properties. In contrast, five isoforms of MreBs in a pathogenic wall-less helical bacterium *Spiroplasma* are involved in its unique motility system driven by a kink propagation along the helical cell. Our integrated biochemical assays show that one isoform of MreBs involved in the swimming of a crustacean pathogen *S. eriocheiris* (SpeMreB1) is exceptionally active in the MreB family proteins and manages the polymerization of another MreB essential for the swimming (SpeMreB5). This study sheds light on an evolutionary mystery how *Spiroplasma* has adapted static MreB proteins to a dynamic phenomenon like its swimming motility.

## Introduction

Motility is the ability of an organism to move directionally from one starting point to another using mechanical energy (converted from chemical energy). Because this ability plays an important role in the survival of organisms, such as obtaining nutrients and evading predators, the evolution of motility has been linked to that of organisms themselves (1). In most bacteria, the peptidoglycan layer, the bacterial cell wall, plays an important role for their motility. The most conserved bacterial motility machineries, flagella and pili (2), require the anchor of the peptidoglycan layer for their propulsion (1). However, because of its conservation across the bacteria kingdom, the peptidoglycan layer is often targeted by natural immunity (3, 4). Therefore, the absence of peptidoglycans is likely advantageous to pathogenic bacteria (1, 5). Certain peptidoglycan-deficient species have evolved unique motility systems that do not rely on conventional bacterial motility machinery requiring the peptidoglycan layer (1).

*Spiroplasma* is a wall-less helical bacterium pathogenic to plants and arthropods. It is often considered industrially problematic, as certain species exhibit strong pathogenicity to host organisms (6–9). *Spiroplasma* swims by propagating its cell helicity switching along the cell axis to generate a propulsive force (10–12). Each *Spiroplasma* species possesses five classes of bacterial actin proteins MreB (SMreB1–5), which contribute to swimming motility (14–18). A recent study suggested that the swimming motility of *Spiroplasma* is related to its pathogenicity (6, 13), as does the motility of other pathogenic organisms (1). Studies of SMreBs are therefore important not only for elucidating the molecular mechanism of *Spiroplasma* swimming, but also for understanding its pathogenic process.

Although MreB is conserved in many bacteria, especially those with elongated cell shapes and cell walls, its role is related to cell shape maintenance rather than directly driving motility (19). In walled-bacteria, MreB filaments bind to the cell membrane and support the formation and localization of a peptidoglycan synthesis complex for the cell elongation (elongasome complex) (20, 21). As peptidoglycan synthesis progress, MreB filaments move along the cell membrane in a direction perpendicular to the major cell axis. In this process, the driving force of MreB movement is the peptidoglycan synthesis rather than the polymerization dynamics of MreB. Thus, MreB filaments are believed to be static during the cell shape maintenance process (22, 23). The static nature of MreB is also suggested by its small conformational change upon polymerization compared to other actin superfamily proteins (24). It is an open question how a dynamic phenomenon like *Spiroplasma* swimming could have emerged in MreB family proteins, which are supposed to be static in other bacterial cells.

Because of its importance in pathogenicity and the evolutionary interest, *Spiroplasma* swimming has been studied from several perspectives, including cellular and molecular biology (25). A previous study using JCVI-syn3B, a minimal synthetic bacterium, revealed that a combination of SMreB5 and either the phylogenetically related SMreB1 or SMreB4 is essential for *Spiroplasma* swimming (15). SMreB5 has been studied biochemically and structurally, and the following properties have been reported (17, 26–28): SMreB5 possesses ATPase and polymerization activities similar to eukaryotic actin; SMreB5 exhibits the polymerization cycle similar to eukaryotic actin, in which pre-ATP hydrolysis and post-phosphate (P_i_) release states are unfavorable for retaining the filaments; SMreB5 forms asymmetric sheets composed of an antiparallel double-stranded filament common to the studied MreBs and parallelly oriented protofilaments; SMreB5 binds to a negatively charged membrane mimicking the *Spiroplasma* cell via its positively charged and unstructured C-terminal tail; and SMreB5 forms bundles depending on its ATPase activity and surface charge. However, SMreB1 and SMreB4 have not been studied because they were not purified in the soluble fraction (27).

In the present study, we solubilized SMreB1 and analyzed it using biochemical techniques. SMreB1 of the crustacean pathogen *Spiroplama eriocheiris* (7) (SpeMreB1) was fused with ProteinS (PrS), which is a spore surface protein of a soil bacterium *Myxococcus xanthus* and has also been reported as a solubilization tag (29). Soluble SpeMreB1 was investigated for its activity, crosstalk with SpeMreB5, and its membrane-binding ability. Based on these analyses, we discuss the functions of SpeMreB1.

## Results

### Preparation of PrS-fused soluble SpeMreB1

As previously reported (27), neither the 6×His-tag fused SpeMreB1 nor SpeMreB4 was soluble (Fig. S1A). This was also the case for SMreB1 and SMreB4, which were derived from different *Spiroplasma* species (Fig. S1B–C, Supplementary data 2). We co-expressed SpeMreB4 and SpeMreB5 because the genes for expressing them are coded tandemly in the genomes of many *Spiroplasma* species (18, 30). However, this approach did not solve the insolubility problem of SpeMreB4 (Fig. S1D). Therefore, we used a solubilization tag for SpeMreB1 and SpeMreB4. We fused two consecutive ProteinS (PrS) molecules at the N-termini of SpeMreB1 and SpeMreB4 (PrS-SpeMreB1 and PrS-SpeMreB4, respectively) and a 6×His-tag at their C-termini (Fig. 1A, Supplementary data 1) and successfully purified them using Ni^2+^-NTA affinity chromatography and gel filtration (Fig. S1E). During gel filtration, all PrS-SpeMreB4 molecules were eluted in the void fraction where the molecular size exceeded the upper separation limit of the column, indicating that PrS-SpeMreB4 formed oligomers. In contrast, PrS-SpeMreB1 was eluted in both the void and monomeric fractions (Fig. 1B). The oligomer formation of PrS-SpeMreB1 and PrS-SpeMreB4 was probably not caused by the soluble aggregation of PrS, as is reported with other solubilization tags (31) because a gel filtration detected the PrS carrying the 6×His-tags at the both N- and C-termini, which was used as the negative control in the following experiments, at the monomeric fraction even at a high concentration for the purification and the following assays (50 μM) (Fig. S2A). In this study, we retained the PrS tag on these SpeMreBs, because the tag removal by factorXa precipitated SpeMreB1 (Fig. S1F).

**Fig. 1.**
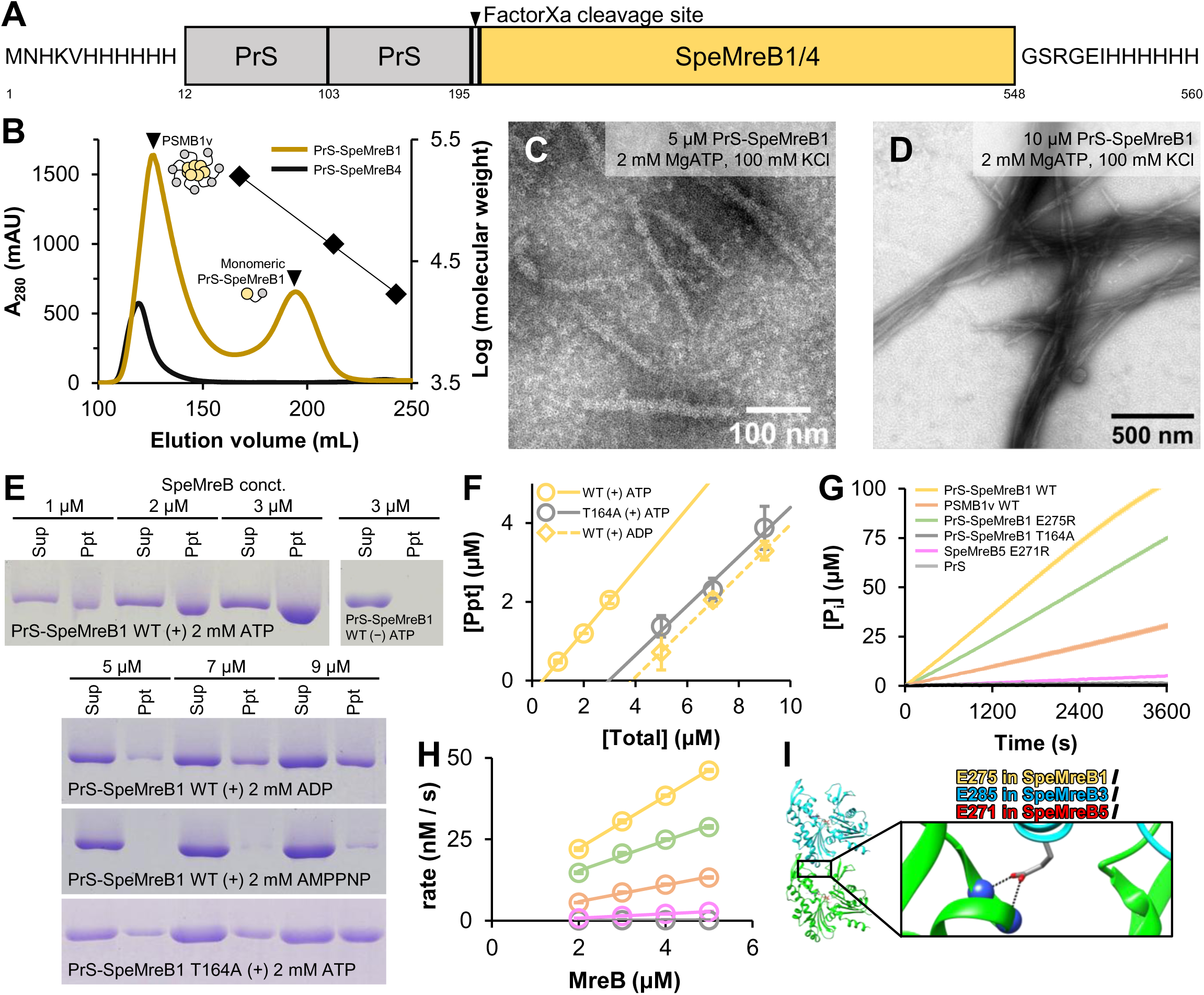
Preparation and evaluation of PrS-SpeMreB1 and PrS-SpeMreB4. (**A**) Domain architecture of PrS-fused SpeMreB1 and SpeMreB4. The N-termini of SpeMreB1 and SpeMreB4 were fused with a 6×His-tag, two consecutive PrS molecules, and a Factor Xa cleavage site. The C-termini of SpeMreB1 and SpeMreB4 were fused with a 6×His-tag. The numbers underneath the figure indicate the residue numbers of PrS-SpeMreB1. (**B**) Gel filtration profiles of PrS-SpeMreB1 WT (dark yellow) and PrS-SpeMreB4 WT (black). Peak positions of PSMB1v WT and the monomeric PrS-SpeMreB1 WT are indicated by closed triangles. Bovine γ-globulin (158 kDa), chicken ovalbumin (44 kDa), and horse myoglobin (17 kDa) were used as the protein size standards, and their elution volumes are indicated by closed diamonds with the linear fit over the log of their molecular weights. (**C–D**) Negative staining EM images of (**C**) 5 μM and (**D**) 10 μM PrS-SpeMreB1 WT in the standard buffer (20 mM Tris-HCl pH 7.5, 100 mM KCl, 5 mM DTT, 2 mM MgCl_2_, 2 mM ATP). (**E**) Sedimentation assays of PrS-SpeMreB1 WT over the nucleotide states and PrS-SpeMreB1 T164A in the presence of ATP. (**F**) Quantification of pellet amounts of PrS-SpeMreB1 WT in the presence of ATP or ADP and PrS-SpeMreB1 T164A in the presence of ATP. The resulting concentrations of the precipitated fractions were plotted against the total SpeMreB concentrations with linear fitting. Error bars indicate the SD of three independent measurements. Critical concentrations were estimated as the *x*-intercept of each linear fit and are summarized in Table 1. (**G–H**) (**G**) Time course and (**H**) concentration dependence of P_i_ release from PrS-SpeMreB1 WT (yellow), PSMB1v WT (orange), PrS-SpeMreB1 E275R (light green), PrS-SpeMreB1 T164A (dark gray), SpeMreB5 E271R (pink), and PrS (light gray). Data are presented as mean ± SD of three independent measurements. In the panel G, protein concentrations were set at 3 μM for SpeMreBs and 5 μM for PrS. P_i_ release rates (k_Pi_) were estimated from the slopes of the linear fits in the panel H and are summarized in Table 2. (**I**) Ribbon representation of two consecutive molecules along the *a*-axis in the crystal of the SpeMreB3-AMPPNP complex (PDB: 7E1G) and the close-up view of its intra-protofilament interaction region. Residue E285 is represented using the stick model with the corresponding residue numbers in SpeMreB1, SpeMreB3, and SpeMreB5.

**Table 1.** Bulk critical concentrations of SpeMreBs estimated from sedimentation assays. Values are indicated as the mean ± SD of three independent measurements, except for PrS-SpeMreB1 WT in the presence of AMPPNP whose critical concentrations could not be determined by the linear fitting due to the small pellet amounts.

| Construct-nucleotide | PrS-SpeMreB1 WT-ATP | PrS-SpeMreB1 WT-ADP | PrS-SpeMreB1 WT-AMPPNP | PrS-SpeMreB1 T164A-ATP | SpeMreB5 WT-ATP | SpeMreB5 WT-ADP | SpeMreB5 WT-AMPPNP | SpeMreB5 T160A-ATP |
| --- | --- | --- | --- | --- | --- | --- | --- | --- |
| C<sub>C</sub> (μM) | 0.41 ± 0.05 | 3.8 ± 0.7 | >= 7 | 3.0 ± 0.2 | 0.14 ± 0.07 | 0.39 ± 0.09 | 0.29 ± 0.11 | 0.15 ± 0.15 |
| Reference | This study | This study | This study | This study | Takahashi D *et al.*, 2022 *Open Biol.* 12(10):220083. | Takahashi D *et al.*, 2022 *Open Biol.* 12(10):220083. | Takahashi D *et al.*, 2022 *Open Biol.* 12(10):220083. | Takahashi D *et al.*, 2022 *Open Biol.* 12(10):220083. |

**Table 2.**
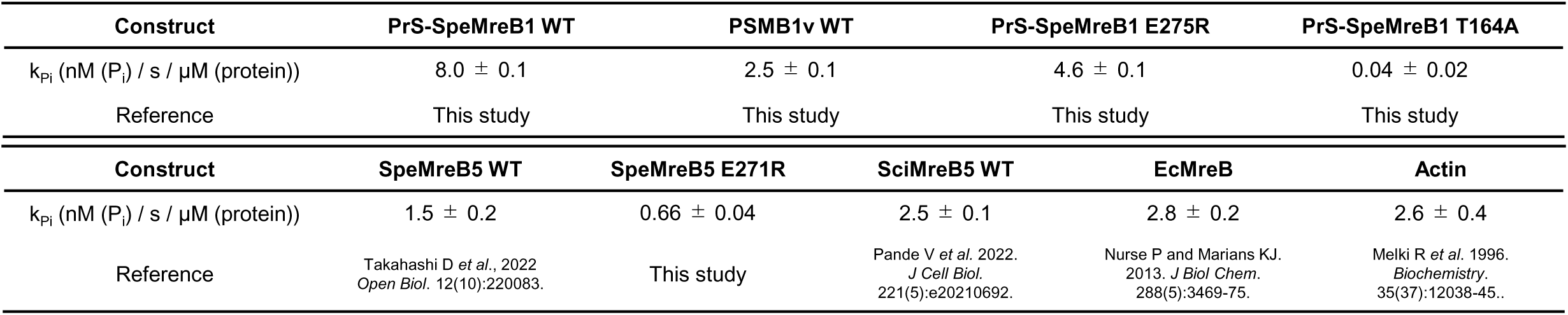
P_i_ release rates of SpeMreBs. Values are indicated as the mean ± SD of three independent measurements.

### Polymerization of SpeMreB1 is highly dynamic among MreB family proteins

We evaluated the assembled structures of PrS-SpeMreB1 after the polymerization of its monomeric fraction by adding ATP. PrS-SpeMreB1 formed sheet structures in the presence of ATP (Fig. 1C). Their 2D averaged image showed the 5.0 ± 0.2 nm subunit repeat consistent with other MreB filaments (17, 20, 27, 28, 32–34), although the protofilament arrangement was obscure probably due to the small particle number available for the image averaging (∼1,000 particles per 50 micrographs) (Fig. S3A). Increasing the PrS-SpeMreB1 concentration from 5 to 10 μM resulted the filament bundling (Fig. 1D), whereas the bundles were partially disassembled by increasing the salt concentration from 100 to 500 mM (Fig. S3B), indicating that the filament bundling of PrS-SpeMreB1 is governed by electrostatic interactions similar to that of SpeMreB5 (26). In contrast, PrS-SpeMreB1 did not polymerize under nucleotide-free (Nf) conditions (Fig. S3C–D).

Next, to evaluate the polymerization activity of PrS-SpeMreB1, we performed sedimentation assays across nucleotide states and estimated each critical concentration (C_C_), which is the minimum concentration required for polymerization (Fig. 1E–F, Table 1). Although PrS-SpeMreB1 did not precipitate under Nf conditions, in the presence of ATP, it precipitated within the C_C_ range compatible with SpeMreB5. In contrast, PrS-SpeMreB1 was less polymerized in the presence of ADP, and its C_C_ was 9.3-fold higher than that in the presence of ATP. This fold change was larger than the C_C_ differences of SpeMreB5 in the presence of ATP and ADP (2.8-fold). PrS-SpeMreB1 hardly polymerized in the presence of AMPPNP (a non-hydrolyzed ATP analog). The C_C_ of an ATPase-depleted variant, PrS-SpeMreB1 T164A, was 7.3-fold higher than that of the wild-type (WT) in the presence of ATP. These results suggest that SpeMreB1 filaments exchange their subunits more actively than SpeMreB5 filaments, depending on the change in the nucleotide state.

Next, to evaluate the ATPase activity of SpeMreB1, we measured P_i_ release. Although P_i_ release was not detected with PrS alone, all SpeMreBs used in this study released P_i_ linearly over time (Fig. 1G). The P_i_ release rate of PrS-SpeMreB1 was 5.3-fold higher than that of SpeMreB5 (27) (Fig. 1H, Table 2) and also 2.9- and 3.1-fold higher than that of *E. coli* MreB, which had the highest record of the P_i_ release rate among the studied MreB family proteins (35) and animal actin (36), respectively. The P_i_ release rate of PrS-SpeMreB1 T164A was 200-fold lower than that of the WT, confirming that SpeMreB1 has an ATP hydrolysis mechanism common to other MreB family proteins, as previously suggested (27). Taken together, these results suggest that the polymerization of SpeMreB1 is highly dynamic among the MreB family proteins.

The linearity of the time-course P_i_ release measurements suggests a futile cycle in which ATP is consumed by SpeMreB monomers, similar to *Geobacillus stearothermophilus* MreB (GsMreB) (32) (Fig. 1G). To confirm this possibility, we created polymerization-deficient variants of SpeMreB1 and SpeMreB5 and measured their P_i_ release rates. Referring to the crystal structure of the SpeMreB3-AMPPNP complex, the E285 residue electrostatically interacts with the positively charged side of the α-helix starting from V216 (27). The corresponding residue of SpeMreB3 E285 is conserved in SMreBs, including SpeMreB1 and SpeMreB5 (E275 and E271 for SpeMreB1 and SpeMreB5, respectively) (16) (Fig. 1I). Therefore, we generated PrS-SpeMreB1 E275R and SpeMreB5 E271R as polymerization-deficient variants. These variants were not polymerized as confirmed by sedimentation assays (Fig. S1G). PrS-SpeMreB1 E275R and SpeMreB5 E271R still exhibited ATPase activities that were 1.7- and 2.3-fold lower, respectively, than those of the corresponding WTs. The decrease in the P_i_ release rate in the polymerization-deficient variant of SpeMreB5 is consistent with the results of a previous study (28). These results indicate that SpeMreB1 and SpeMreB5 also possess futile cycles similar to those of GsMreB (32) and that these ATPase activities are promoted by polymerization.

In addition to SpeMreB1, we performed sedimentation assays of PrS-SpeMreB4 eluted in the void fraction (Fig. 1B). Although PrS-SpeMreB4 precipitated regardless of ATP (Fig. S1H), filamentous structures were not formed, even in the presence of ATP (Fig. S3E), suggesting that PrS-SpeMreB4 precipitation in the sedimentation assays was caused by PrS-SpeMreB4 aggregation rather than polymerization. Therefore, we excluded PrS-SpeMreB4 from further analyses.

### Polymerization of SpeMreB5 is required for the binding between SpeMreB1 and SpeMreB5

Next, we evaluated the interactions between SpeMreB1 and SpeMreB5. However, under mixed conditions of monomeric WTs, it was difficult to distinguish whether the obtained signals were derived from their interactions or promoted polymerization. Therefore, we used polymerization-deficient SpeMreB1 variants to detect their interactions with SpeMreB5. In addition to the PrS-SpeMreB1 E275R variant, we analyzed the interaction of PrS-SpeMreB1 WT eluted in the void fraction at gel filtration (PSMB1v) (Fig. 1B). The size distribution of PSMB1v did not depend on the incubation and the presence of ATP (Fig. S2B–E), and it was less precipitated by an ultracentrifugation even in the presence of ATP (Fig. S2F). These results suggest that the subunits within the PSMB1v oligomers are not exchanged with other PrS-SpeMreB1 molecules. Nevertheless, PSMB1v retained several characteristics of the PrS-SpeMreB1 monomer as follows: the presence of the ATPase activity, although its P_i_ release rate was 3.2-fold lower than that of the PrS-SpeMreB1 monomer (Table2, Fig. 1G–H); and its secondary structural content, which was consistent with that of PrS-SpeMreB1 as estimated by circular dichroism (CD) measurements (Table 3, Fig. S2G). Therefore, we used PSMB1v to analyze its interaction with SpeMreB5.

**Table 3.**
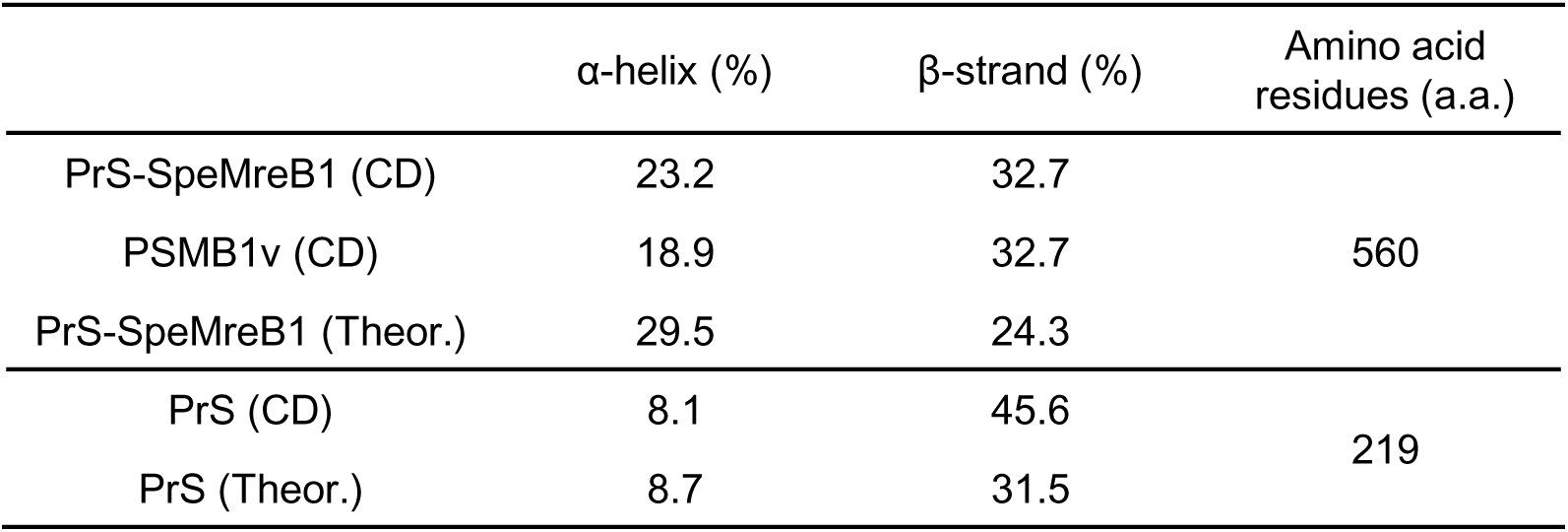
Secondary structure contents and number of amino acid residues in PrS-SpeMreB1 and PrS. The secondary structure contents in the rows labeled CD and Theor. were estimated from the CD spectra (Fig. S2G) and amino acid sequences, respectively.

Both PSMB1v and PrS-SpeMreB1 E275R were precipitated by ultracentrifugation in the presence of ATP-polymerized SpeMreB5 (Fig. 2A–B), although PrS itself did not precipitate, even in the presence of SpeMreB5 and/or ATP (Fig. S4A). These results indicate that SpeMreB1 binds to SpeMreB5 filaments, which is probably not the artefact by the formation of PSMB1v or the E275R mutation. Deletion of the C-terminal nine SpeMreB5 residues involved in the membrane binding and nucleating the bundle formation (SpeMreB5 ΔC9 variant) (26, 28) did not affect to its affinity for PSMB1v (Fig. 2A second top gel, 2B).

**Fig. 2.**
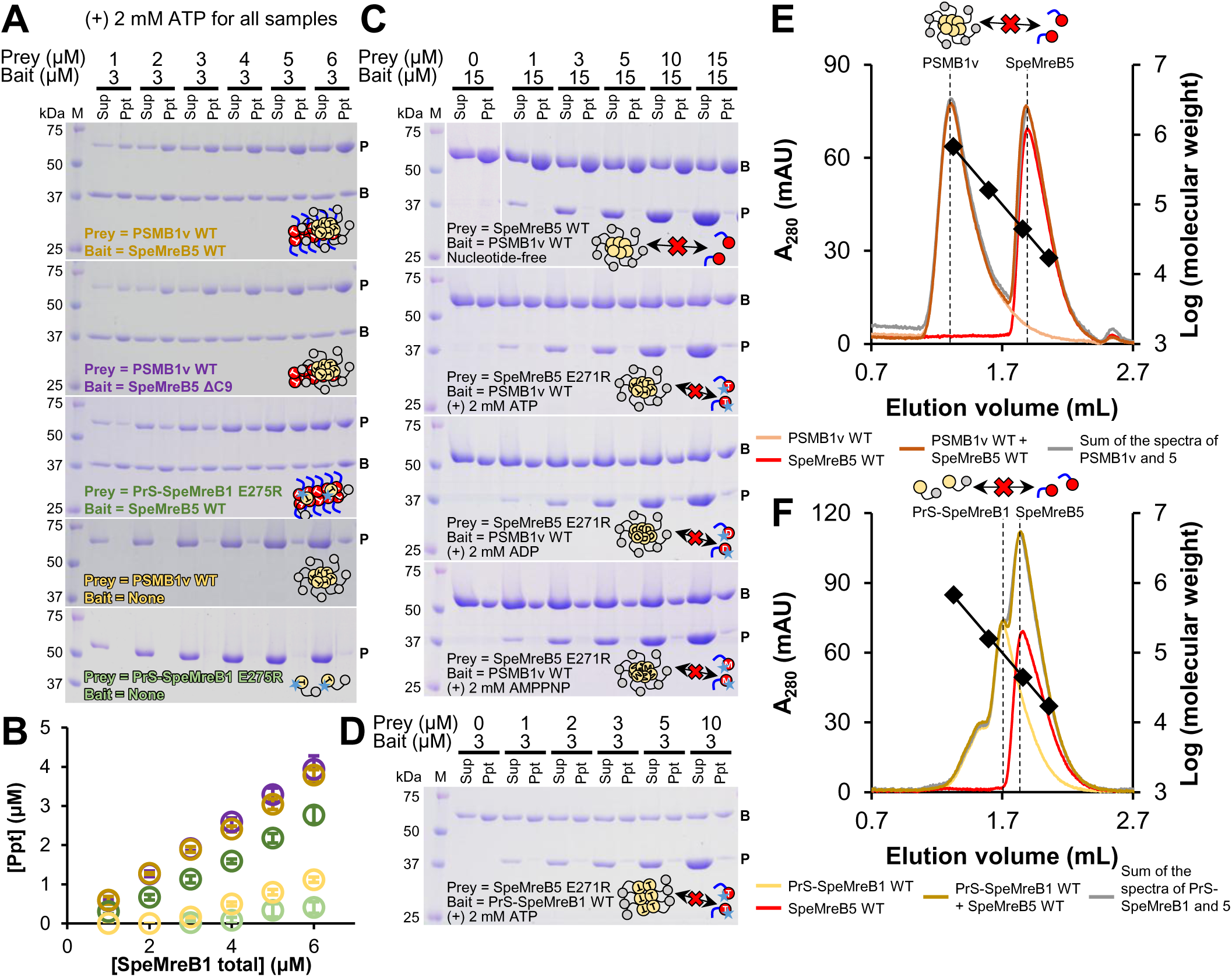
Interaction between SpeMreB1 and SpeMreB5. For co-sedimentation assays, protein size standards were visualized in lane M with the molecular masses of each band on the left side. The band positions of prey and bait in the gel images are indicated by P and B, respectively. The result scheme of each experiment is shown as a cartoon reflecting the assembly states (monomeric, polymerized, or oligomerized), nucleotide states (empty, T, D, and M symbols for Nf, ATP, ADP, and AMPPNP, respectively), and mutations (star for the polymerization-deficient mutation and the lack of blue line for the C-terminus deletion of SpeMreB5) of PrS-SpeMreB1 (yellow circle connected with a gray circle indicating PrS) and SpeMreB5 (red circle with the C-terminal region as a blue line). (**A**) Co-sedimentation assays of PrS-SpeMreB1 and SpeMreB5 variants in the pairs indicated in each gel image in the presence of 2 mM ATP. The concentration of SpeMreB5 variants was constant at 3 μM, whereas that of PrS-SpeMreB1 variants varied from 1 μM to 6 μM. (**B**) Quantification of pellet amounts of PrS-SpeMreB1 variants in the co-sedimentation assays with the 3 μM SpeMreB5 variant. The resulting concentrations of the precipitated fractions were plotted against the total concentrations by open circles with the matched color scale to Fig. 2A. Error bars indicate the SD of three independent measurements. (**C**) Co-sedimentation assays of PSMB1v WT and SpeMreB5 WT in the Nf condition or SpeMreB5 E271R in the presence of 2 mM ATP, ADP, or AMPPNP. The concentration of PSMB1v WT was constant at 15 μM, whereas that of SpeMreB5 variants varied from 0 μM to 15 μM. The centrifugation time was set to 30 min, which was short enough to minimize the precipitation of the high concentration of SpeMreB5 monomers. (**D**) Co-sedimentation assay of 3 μM PrS-SpeMreB1 WT and 0 μM to 10 μM SpeMreB5 E271R in the presence of 2 mM ATP. (**E–F**) Size exclusion chromatography of (**E**) PSMB1v WT and (**F**) PrS-SpeMreB1 WT with SpeMreB5. Peak positions of PSMB1v WT, PrS-SpeMreB1 WT, and SpeMreB5 WT are indicated by dashed lines. Bovine thyroglobulin (670 kDa), bovine γ-globulin (158 kDa), chicken ovalbumin (44 kDa), and horse myoglobin (17 kDa) were used for the protein size standards, and their elution volumes are plotted by closed diamonds with the linear fit over the log of their molecular weights. (**E**) Gel filtration spectra of 10 μM PSMB1v WT (light orange), 10 μM SpeMreB5 WT (red), the mixture of 10 μM PSMB1v WT and 10 μM SpeMreB5 WT (brown), and the sum of spectra from independent loads of 10 μM PSMB1v WT and 10 μM SpeMreB5 WT (gray). (**F**) Gel filtration spectra of 10 μM PrS-SpeMreB1 WT (yellow), 10 μM SpeMreB5 WT (red), the mixture of 10 μM PrS-SpeMreB1 WT and 10 μM SpeMreB5 WT (dark yellow), and the sum of spectra from independent loads of 10 μM PrS-SpeMreB1 WT and 10 μM SpeMreB5 WT (gray).

Next, we attempted to detect the interaction between SpeMreB5 monomers and SpeMreB1. Due to the large size of PSMB1v, it exhibited noticeable precipitation at a high concentration. Taking advantage of this property, we performed co-sedimentation assays of PSMB1v and monomeric SpeMreB5 WT under Nf conditions or SpeMreB5 E271R over the nucleotide states (Fig. 2C). Most SpeMreB5 molecules remained in the supernatant under all conditions. In addition, we performed co-sedimentation assays with PrS-SpeMreB1 WT and SpeMreB5 E271R in the presence of ATP to study the interaction between polymerized SpeMreB1 and SpeMreB5 monomers (Fig. 2D). Most SpeMreB5 E271R molecules remained in the supernatant, whereas PrS-SpeMreB1 precipitated. The interaction between the SpeMreB5 monomer and SpeMreB1 variants was further assessed using size-exclusion chromatography (Fig. 2E–F). The peak positions and magnitudes did not change in the mixed conditions of PSMB1v or PrS-SpeMreB1 and SpeMreB5 compared with when each was independently loaded. These results indicate that the polymerized form of SpeMreB5 is essential for the interaction between SpeMreB1 and SpeMreB5.

### SpeMreB1 decreases SpeMreB5 filament amount depending on the nucleotide states

A previous study demonstrated that SMreB1 expressed in *E. coli* cells inhibited the filament formation of SMreB2, which was co-expressed as a model of a phylogenetic group including SMreB5 (37). Inspired by this study, we evaluated the effects of SpeMreB1 on SpeMreB5 filaments using co-sedimentation assays in the presence of different nucleotides (Fig. 3A–B, S4B–D). In the presence of ATP, the amount of SpeMreB5 precipitate did not change when mixed with approximately thrice higher concentrations of PrS-SpeMreB1 or PSMB1v.

**Fig. 3.**
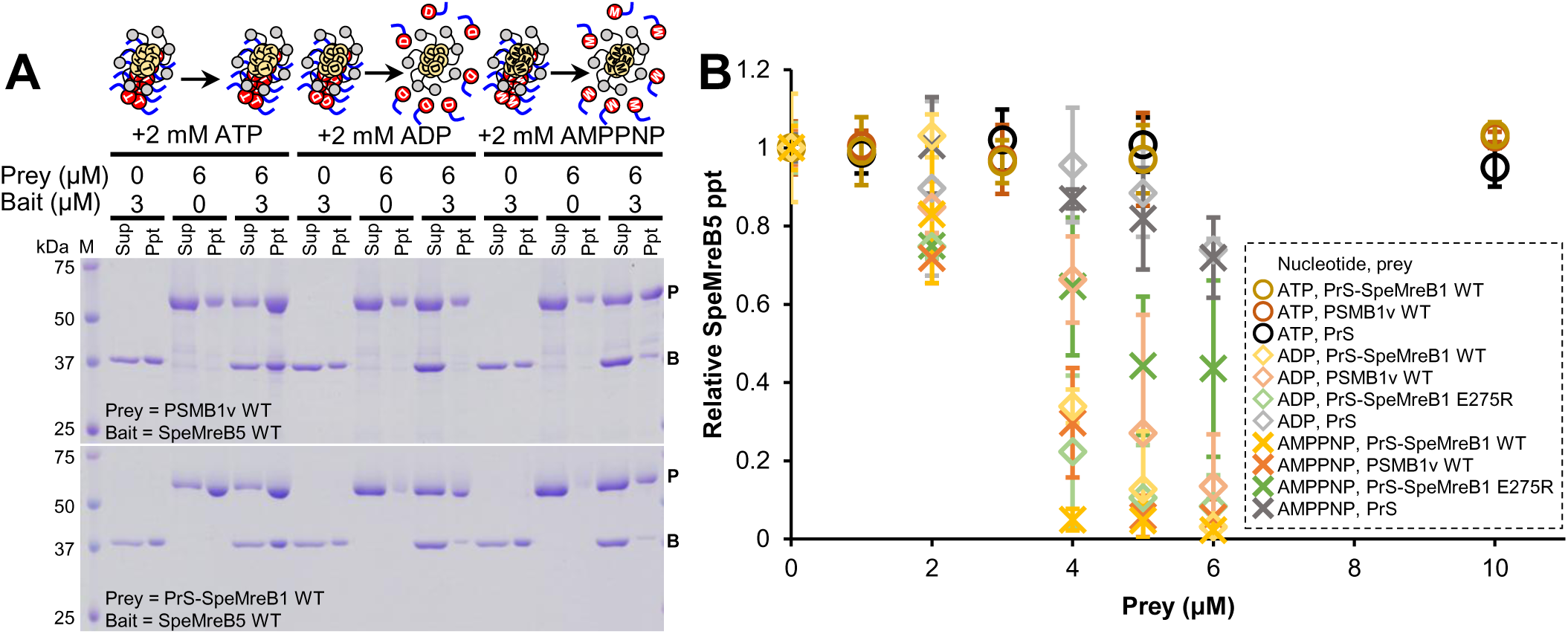
Effects of SpeMreB1 on SpeMreB5. (**A**) Co-sedimentation assays of 3 μM SpeMreB5 WT and 6 μM PSMB1v WT (top) or PrS-SpeMreB1 WT (bottom) in the presence of 2 mM ATP, ADP, or AMPPNP. Protein size standards were visualized in lane M with the molecular masses of each band on the left side. The band positions of prey and bait in the gel images are indicated by P and B, respectively. The result scheme of each co-sedimentation assay of PSMB1v and SpeMreB5 is shown as a cartoon reflecting the assembly states (monomeric, polymerized, or oligomerized) and nucleotide states (empty, T, D, and M symbols for nucleotide-free, ATP, ADP, and AMPPNP, respectively) of PrS-SpeMreB1 (yellow circle connected with gray circle indicating PrS) and SpeMreB5 (red circle with the C-terminal region as a blue line). (**B**) Relative pellet amounts of SpeMreB5 WT in co-sedimentation assays with an SpeMreB1 variant and a nucleotide. The total SpeMreB5 WT concentration was constant at 3 μM, whereas that of PrS-SpeMreB1 variants varied from 0 μM to 10 μM as shown in Fig. S4B–D. The resulting concentrations of the precipitated fractions were plotted against the total concentrations of an SpeMreB1 variant. The colors of the plots indicate the proteins added versus SpeMreB5 WT (yellow-, orange-, green-, and gray-scaled colors for PrS-SpeMreB1 WT, PSMB1v WT, PrS-SpeMreB1 E275R, and PrS, respectively). The shapes of the plots indicate the added nucleotide (open circles, open diamonds, and crosses for 2 mM ATP, ADP, and AMPPNP, respectively). Error bars indicate the SD from three independent measurements.

Intriguingly, in the presence of ADP or AMPPNP, the amount of SpeMreB5 precipitate decreased in the presence of PrS-SpeMreB1 or PSMB1v. In these nucleotide states, SpeMreB5 mostly did not precipitate in the presence of twice the concentration of PrS-SpeMreB1 or PSMB1v. This phenomenon also occurred with PrS-SpeMreB1 E275R in the presence of ADP (in the presence of AMPPNP and PrS-SpeMreB1 E275R, the precipitation of SpeMreB5 did not decrease below 55%). In contrast, the precipitate decrease in SpeMreB5 by PrS under conditions corresponding to SpeMreB1 variants remained at 26.1% and 28.1% in the presence of ADP and AMPPNP, respectively. These results suggest that SpeMreB1 dissociated SpeMreB5 filaments.

### SpeMreB1 polymerization is essential for *Spiroplasma* swimming

The above experiments revealed that the polymerization-deficient variant of SpeMreB1 partially retained the ATPase activity, interaction with SpeMreB5 filaments, and the ability to decrease SpeMreB5 filaments (Fig. 1G–H, 2A–B, 3A–B). To examine the significance of SpeMreB1 polymerization in *Spiroplasma* swimming, we investigated the swimming behavior of cells expressing an SpeMreB1 variant with SpeMreB5 WT using the reconstitution system of the minimal synthetic bacterium JCVI-syn3B (15).

Although syn3B cells co-expressing SpeMreB1 WT and SpeMreB5 WT were motile, as observed in *Spiroplasma*, those co-expressing SpeMreB1 E275R and SpeMreB5 WT were not (Fig. 4A–B). Protein profile analysis revealed that bands corresponding to the SpeMreB1 variant and SpeMreB5 WT were detected in both strains (Fig. 4C, Table S1), suggesting that the loss of swimming motility in the cells co-expressing SpeMreB1 E275R and SpeMreB5 WT was not caused by an aberrant SpeMreB expression. These results indicate that SpeMreB1 polymerization is essential for *Spiroplasma* swimming.

**Fig. 4.**
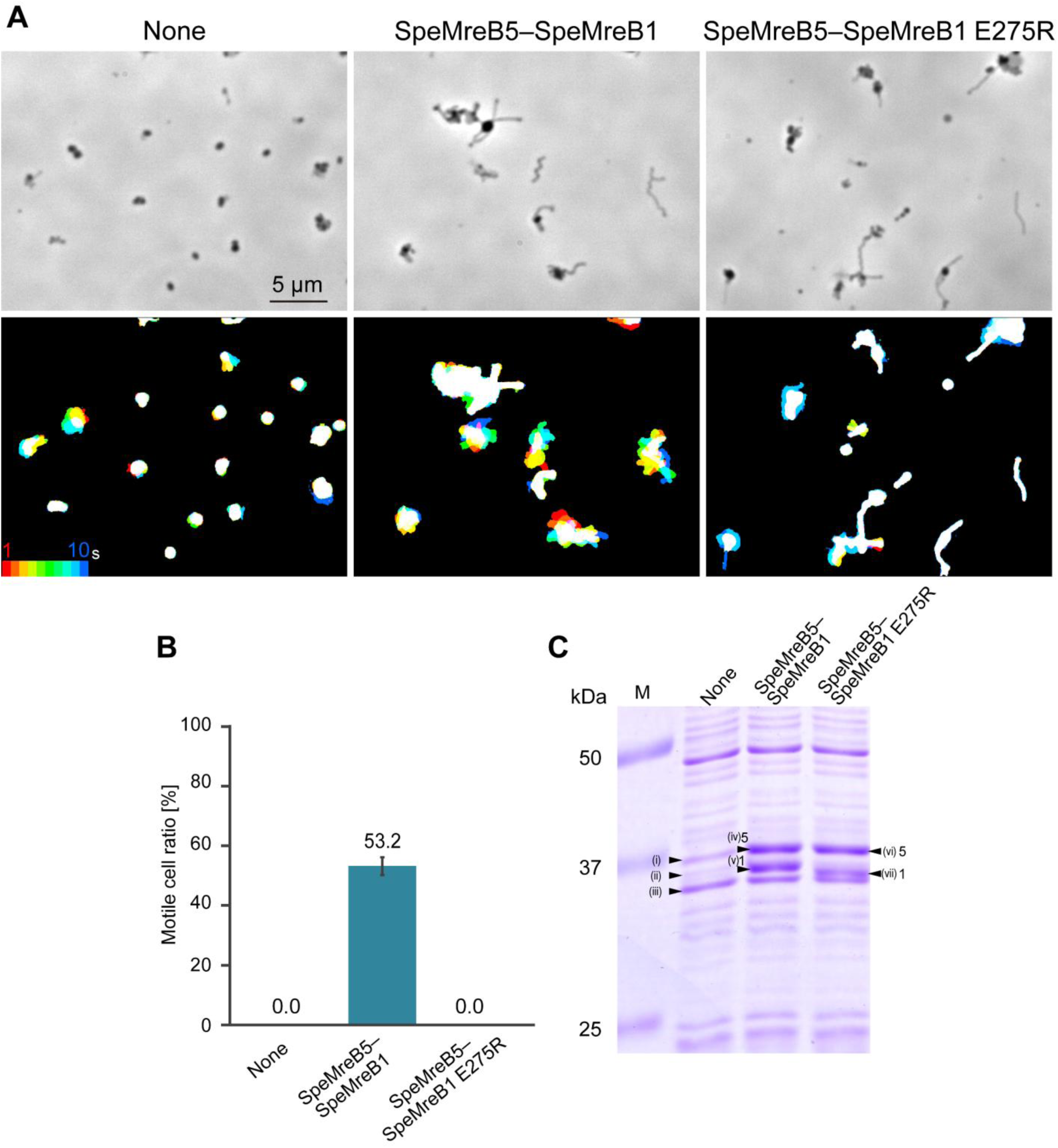
Behavior of syn3B cells expressing an SpeMreB1 variant and SpeMreB5 WT. (**A**) (Upper) Phase-contrast field images of control syn3B cells carrying the antibiotic selection maker (left) and those co-expressing SpeMreB1 WT and SpeMreB5 WT (middle) and SpeMreB1 E275R and SpeMreB5 WT (right). (Lower) Trajectories of cells corresponding to each upper image for 10 s. Video frames were clipped for every 1 s, colored from red to blue in a rainbow-colored manner, and superimposed onto one image. (**B**) The motile cell ratio of the syn3B expressing SpeMreBs analyzed for three individual fields (110.8 × 62.3 μms) including 911, 545, and 563 cells, respectively. (**C**) Protein profiles of syn3B strains. Protein size standards are shown in lane M with the molecular masses of each band on the left side. The marked protein bands were identified by peptide mass fingerprinting as summarized in Table S1. The bands of SpeMreB1 and SpeMreB5 are marked as 1 and 5, respectively.

### SpeMreB1 binds to negatively charged lipids in any nucleotide states

As the motility machinery of *Spiroplasma* is present underneath the cell membrane (14, 38–40), it is important to study the membrane-binding ability of each component of the motility machinery. Although the membrane binding of SMreB1 is largely unclear, its binding to negatively charged lipids has been experimentally demonstrated for SMreB5 (28). Therefore, we investigated the membrane binding of PrS-SpeMreB1 based on the co-sedimentation assay with liposomes. In this assay, we reduced the centrifugation speed (from 436,000 ×*g* to 100,000 ×*g*) and time (from 90 min to 30 min) in the sedimentation assays without liposomes to reduce the background precipitation of SpeMreB filaments. PrS-SpeMreB1 monomers co-precipitated with liposomes with a lipid composition mimicking that of the *Spiroplasma* cell membrane (SpiroLipid liposome), whereas PrS did not (Fig. 5A), indicating that SpeMreB1 binds to the membrane, contrary to our previous prediction based on the absence of membrane-binding sequences common to MreB family proteins (16). The precipitated fractions of PrS-SpeMreB1 and SpeMreB5 were 34.9% and 51.4%, respectively (Fig. 5B), at the saturated liposome concentration, indicating that the membrane binding ability of SpeMreB1 is slightly weaker than that of SpeMreB5.

**Fig. 5.**
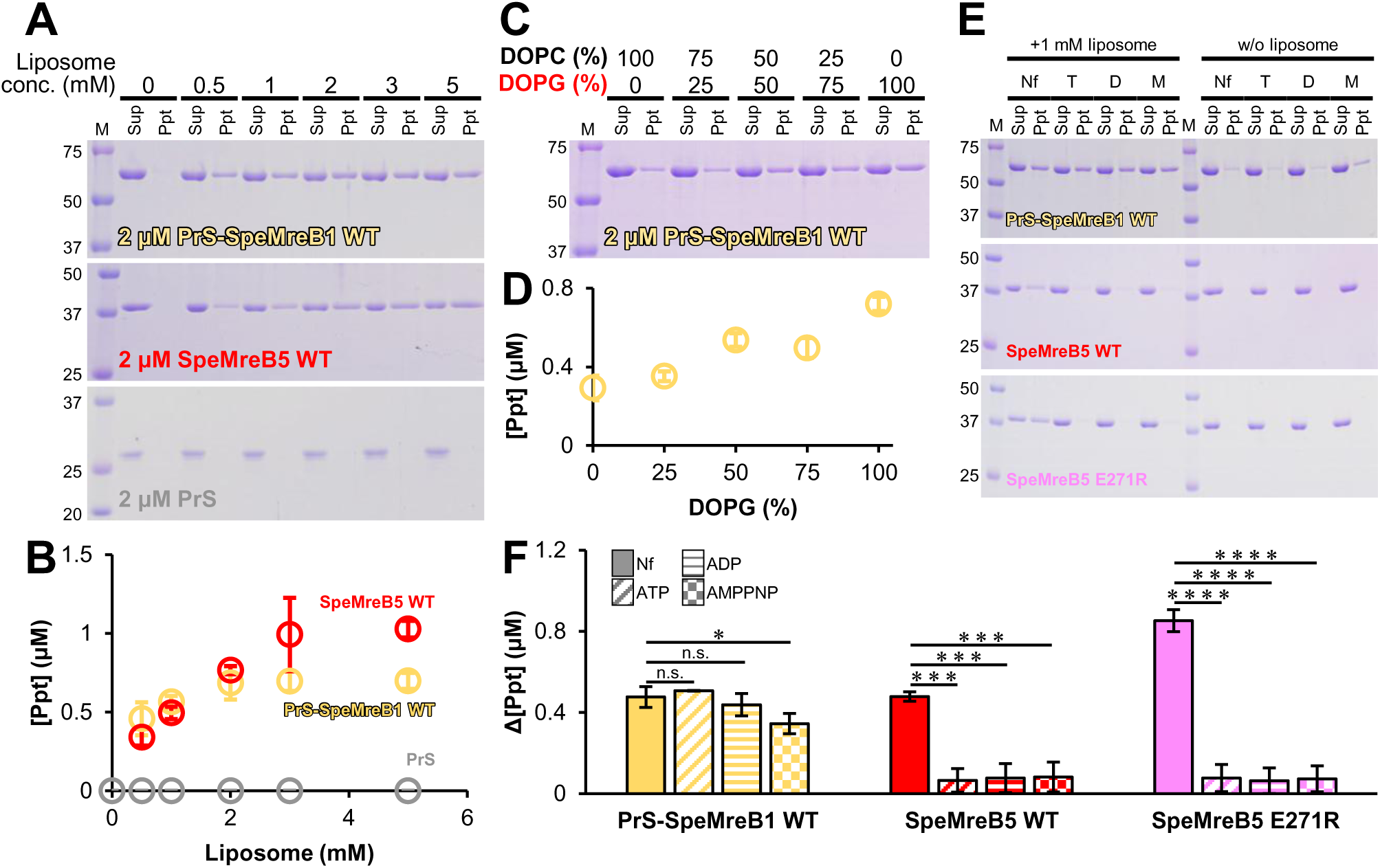
Liposome binding of SpeMreBs. For gel images, protein size standards were visualized in lane M with the molecular masses of each band on the left side. (**A**) Liposome binding assays of 2 μM PrS-SpeMreB1 WT (top), 2 μM SpeMreB5 WT (middle), and 2 μM PrS (bottom) for concentrations of SpiroLipid liposomes. (**B**) Quantification of pellet amounts in the liposome binding assays. The resulting concentrations of the precipitated fractions were plotted over the total liposome concentrations by yellow (2 μM PrS-SpeMreB1 WT), red (2 μM SpeMreB5 WT), and gray (2 μM PrS) open circles. Error bars indicate the SD of three independent measurements. (**C**) Liposome binding assay of 2 μM PrS-SpeMreB1 WT in the presence of 1 mM liposomes with different DOPC and DOPG ratios shown at the top of each lane. DOPG is labeled in red characters as indicative of its negative charge. (**D**) Quantification of pellet amounts in the liposome binding assays of 2 μM PrS-SpeMreB1 WT in the presence of 1 mM liposomes with different DOPC and DOPG ratios plotted against the DOPG ratio. Error bars indicate the SD of three independent measurements. (**E**) Liposome binding assays of 2 μM PrS-SpeMreB1 WT (top), 2 μM SpeMreB5 WT (middle), and 2 μM SpeMreB5 E271R (bottom) in the presence of 1 mM SpiroLipid liposome and 1 mM nucleotide as indicated at the top of each lane (Nf, T, D, M for nucleotide-free, with ATP, with ADP, and with AMPPNP, respectively). (**F**) Quantification of pellet amounts in the liposome binding assays in the presence of 1 mM SpiroLipid liposome and 1 mM nucleotide. Precipitate concentration differences in the presence and absence of liposomes were summarized for each construct. The patterns in the bars indicate the added nucleotides (solid, diagonal stripe, horizontal stripe, and check patterns for nucleotide-free, with ATP, with ADP, and with AMPPNP, respectively). Error bars indicate the SD of three independent measurements. Symbols indicate *p*-values supported by Student’s *t*-test (* *p* < 0.05, *** *p* < 1.0 × 10^−3^, **** *p* < 1.0 × 10^−4^, and n.s. *p* > 0.05).

To identify the lipid molecules involved in the binding to SpeMreB1, we performed liposome binding assays of PrS-SpeMreB1 using liposomes composed of one of the following four lipids in the SpiroLipid liposome: (1,2-dioleoyl-sn-glycero-3-phospho-(1’-rac-glycerol) [DOPG], 1,2-dioleoyl-sn-glycero-3-phosphocholine [DOPC], sphingomyelin [SM], and cardiolipin [CL]) (Fig. S4E). PrS-SpeMreB1 bound to all four liposomes but strongly bound to those composed of DOPG or CL, both of which are negatively charged. To further examine the negative charge dependence of lipids for binding with SpeMreB1, we performed liposome-binding assays of PrS-SpeMreB1 using liposomes with different ratios of DOPG and DOPC. As the ratio of DOPG increased, the affinity between PrS-SpeMreB1 and the liposomes became high (Fig. 5C–D). These results indicate that SpeMreB1 binds to the membrane including negatively charged lipids like SMreB5 (28). To confirm the consistency of the negatively charged lipid dependence of the membrane binding of SpeMreB1, we modeled the structure of full-length SpeMreB1 using AlphaFold2 (41) (Fig. S4F). MreB is composed of four subdomains (IA, IB, IIA, and IIB) (17, 20, 27, 28, 32, 34), in which the side of the subdomains IA and IB are involved in the membrane binding of all studied MreBs (21, 28). SpeMreB1 possesses a positively charged region throughout its IA and IB subdomains, allowing it to bind to the membrane.

Next, we evaluated the membrane binding of SpeMreB1 to liposomes in the presence of nucleotides (Fig. 5E–F). Even in the presence of nucleotides, PrS-SpeMreB1 and SpeMreB5 precipitate poorly at a reduced centrifugation force in the absence of liposomes. SpeMreB5 did not bind to the liposomes in the presence of nucleotides, which is consistent with a previous study (28). This phenomenon was independent of polymerization, as was confirmed for SpeMreB5 E271R. In contrast, the co-precipitate amounts of PrS-SpeMreB1 with liposomes were less affected than that of SpeMreB5. These results indicate that the membrane-binding region of SpeMreB1 is still functional after binding to a nucleotide, unlike SpeMreB5.

## Discussion

In this study, we solubilized SpeMreB1 by fusing it with PrS and evaluated its activity, crosstalk with SpeMreB5, and membrane binding. The β-strand content of PrS-SpeMreB1 used in this study was 8.4% higher than that predicted from its amino acid sequence (Table 3). Considering that the β-strand content of PrS, which occupies 39.1% of the residues in PrS-SpeMreB1, estimated from CD measurements was 14.1% higher than that predicted from the sequence, 5.5% of the 8.4% difference could be explained (14.1% × 0.391 = 5.5%). Excluding this effect, PrS-SpeMreB1 probably adopted the structure as predicted because the differences in α-helix and β-strand content of PrS-SpeMreB1 in experiments and prediction remained at 6.3% and 2.9%, respectively. The α-helix content differences between PrS-SpeMreB1 and PSMB1v (4.3% lower in PSMB1v than in PrS-SpeMreB1) may reflect partial denaturation of PSMB1v (Table 3). However, this structural difference had little effect on the characteristics of SpeMreB1 as PSMB1v retained the characteristic activity of PrS-SpeMreB1 (Figs. 1, 3, and 5).

Simultaneously with nucleotide-dependent polymerization (Fig. 1C–F), we found the highest fold-changes in C_C_ in nucleotide states and P_i_ release rates of PrS-SpeMreB1 in the MreB family proteins (27, 28, 32, 33, 35, 36, 42–45) (Tables 1 and 2). These findings suggest that SpeMreB1 is exceptionally dynamic in the MreB family proteins. A previous study demonstrated that SMreB1 filaments formed in *E. coli* cells were static (37). Our finding of SpeMreB1 bundles (Fig. 1D) suggests that the filaments observed in *E. coli* cells were in the form of bundles. Based on experiments using polymerization-deficient variants (Fig. 1G–I), we confirmed the ATPase futile cycle in PrS-SpeMreB1 and SpeMreB5, as monitored in GsMreB (32). Time-course P_i_ release measurements of PrS-SpeMreB1 E275R and SpeMreB5 E271R revealed that the P_i_ concentration increased proportionally with time (Fig. 1G). Based on these findings, we propose an updated SpeMreB polymerization cycle (Fig. 6A). First, SpeMreBs can hydrolyze ATP even in the monomeric state, with P_i_ release likely being the rate-limiting step. Monomeric SpeMreBs in either ATP or ADP-P_i_ states are most favorable for initiating polymerization, leading to accelerated P_i_ release. In the case of SpeMreB1, most monomers probably polymerize in the ADP-P_i_ state, given their low polymerization ability in the presence of AMPPNP (Fig. 1E). In contrast, SpeMreB5 displayed only two-fold differences in C_C_ in the presence of AMPPNP and ATP (27) (Table 1) and slow P_i_ release in the polymerization-deficient variant (Table 2, Fig. 1H), suggesting that SpeMreB5 can polymerize before hydrolyzing ATP. In this reaction path, SpeMreB5 follows a polymerization cycle without considering the ATPase cycle in the monomeric state, as previously described (27).

**Fig. 6.**
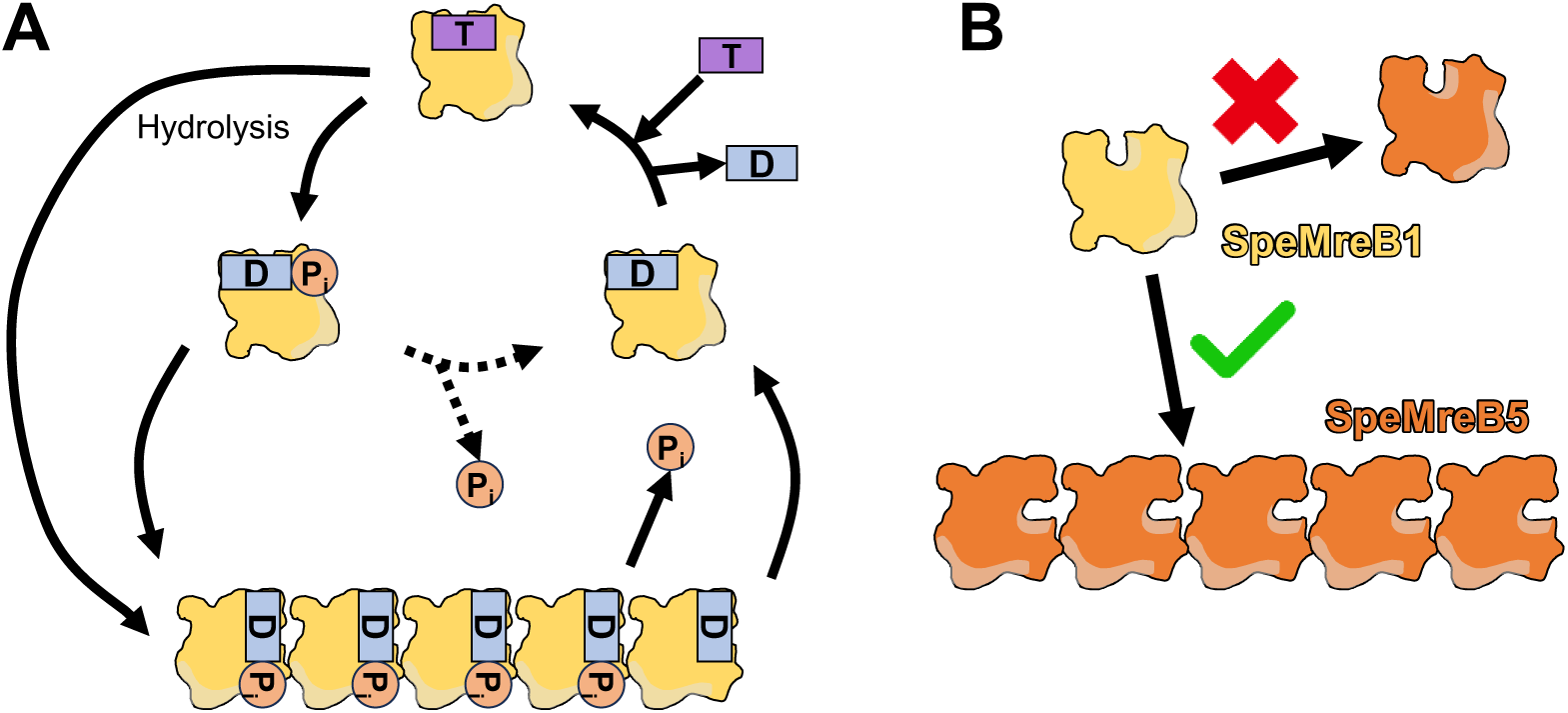
Working models and summary of SpeMreB1 molecular features. (**A**) Working model of the polymerization dynamics of SpeMreB1. The monomeric SpeMreB1 hydrolyzes ATP with the rate-limiting step of P_i_ release. During this cycle, the monomer in the ATP and/or ADP-P_i_ state enters the polymerization cycle, resulting in a higher P_i_ release rate. (**B**) Mode of SpeMreB1 binding to SpeMreB5 filaments, but not to SpeMreB5 monomers.

Polymerization of SpeMreB5 is essential for the interaction between SpeMreB1 and SpeMreB5 (Fig. 2, 6B). In addition, we found a decrease in SpeMreB5 filaments in the presence of SpeMreB1, depending on the nucleotide state (Fig. 3A–B). This result is consistent with a previous study, in which the filament formation of SMreB2, the most identical SMreB to SMreB5, in *E. coli* cells was inhibited by the co-expression of SMreB1 (37). This phenomenon was probably not caused by either the sequestration of SpeMreB5 monomers by SpeMreB1 (Fig. 3B), nor by co-polymerization of SpeMreB1 and SpeMreB5 (Fig. 2A–B). Assuming that MreB5 filament destabilization is involved in the force generation cycle of *Spiroplasma* swimming, it is plausible that SpeMreB1, rather than inhibiting polymerization, destabilizes SpeMreB5 filaments, which may be essential to prevent the polymerization dynamics from stalling. Polymerization of SpeMreB1 was essential for *Spiroplasma* swimming (Fig. 4), suggesting that the interaction between polymerized SpeMreB1 and SpeMreB5 is essential in force generation for this motility. Although the decrease in SpeMreB5 filaments by SpeMreB1 depended on their nucleotide states, the atomic structures of SMreB5 were not substantially different between the ADP- and AMPPNP-bound states (28). Therefore, the decrease in the SpeMreB5 filament amount may be due to the differences of its mechanical properties across the nucleotide state. A previous study has reported that actin filaments become flexible upon P_i_ release, whereas their structure remains largely unchanged. The change in the mechanical properties of actin filaments has been thought to affect the specificity of actin-binding proteins, such as cofilin, which selectively severs actin filaments in the ADP-bound state (46).

We found that SpeMreB1 bound to a membrane containing negatively charged lipids (Fig. 5). In walled bacteria, MreB binds to the membrane via an amphipathic helix at the N-terminus and/or consecutive hydrophobic residues in the hydrophobic loop region in subdomain IA (21). Because most SMreB1s and SMreB4s do not possess these sequences, we predicted that SMreB1 and SMreB4 would not bind to the membrane (16). However, SpeMreB1 bound to the membrane even without the membrane-binding regions common to walled-bacterial MreB (Fig. 5). This finding suggests that SpeMreB1 functions underneath the cell membrane. In addition, SMreB5 binds to the membrane via its positively charged C-terminal region, which has not been reported in the MreBs of walled bacteria (28). These findings suggest that SMreB1 and SMreB5 have evolved unique membrane-binding mechanisms for driving *Spiroplasma* swimming. In this study, we confirmed the dissociation of SMreB5 from the membrane in the presence of a nucleotide (Fig. 5E–F), although independently expressed SMreB1 and SMreB5 in *Mycoplasma mycoides* localized underneath the cell membrane as filaments (14). This inconsistency suggests that other factors that are not reproducible in *in vitro* liposome-binding assays, such as membrane curvature and the excluded volume effect, may control the membrane binding of SMreB filaments.

In conclusion, we demonstrated the dynamic properties, membrane-binding ability, and crosstalk with SpeMreB5 filaments of SpeMreB1. These findings suggest that SpeMreB1 functions as a “molecular motor” to exert the force on a “cytoskeleton” composed of relatively static SpeMreB5 filaments to propel *Spiroplasma* swimming.

## Materials and Methods

### Phylogenetic analysis

The phylogenetic tree (Fig. S1B) was constructed using the maximum likelihood method based on all SMreB1 and SMreB4 sequences reported on March 3, 2021, as previously described (16). Bootstrap supports were estimated from 1,000 alignment samples. Ancestor estimation was performed using MEGA-X, as described previously (16, 47).

### Cloning and expression of SpeMreB

Constructs for the expression of SpeMreB1, SpeMreB4, and SpeMreB5 fused with a 6× His-tag were prepared as the fusions of these genes with codon optimization for *E. coli* expression and a pCold-15b vector, as described in our previous study (27). Expression plasmids for the other SMreB1s and SMreB4s were constructed as fusions of pCold-15b as described previously (27). The plasmid for co-expressing SpeMreB4 and SpeMreB5 was constructed by inserting the codon-optimized *spemreB5* gene with the 5’-extension of GGATCCTAATTTTGTTTAACTTTAAGAAGGAGATAT, which carries an *E. coli* ribosome-binding site to the downstream of the codon-optimized *spemreB4* gene in pCold-15b. To construct expression plasmids for PrS-SpeMreB1 and PrS-SpeMreB4, the codon-optimized genes for *E. coli* expression were excised using NdeI and BamHI restriction enzymes and inserted into pCold-PrS (provided by Dr. Yoshihiro Yamaguchi, Osaka Metropolitan University, Japan), in which two consecutive PrS molecules are fused to the N-terminus of the protein of interest (Supplementary Data 1) and expressed by the pCold expression system with the selective marker of ampicillin. These SpeMreBs carry the N- and C-terminal extensions MNHKVHHHHHHMANITVFYNEDFQGKQVDLPPGNYTRAQLAALGIENNTISSVKVPPGVKAILYQNDGFAGDQIEVVANAEELGPLNNNVSSIRVISVPVQPRMANITVFYNEDFQGKQVDLPPGNYTRAQLAALGIENNTISSVKVPPGVKAILYQNDGFAGDQIEVVANAEELGPLNNNVSSIRVISVPVQPRGTIEGRH and GSRGEIHHHHHH, respectively. An empty pCold-PrS vector was used to express PrS. *Thermotoga maritima* MreB (TmMreB) was constructed as a fusion with an 8× His-tag by inserting the codon optimized sequence to pSY5 vector (48). Each construct was transformed into *E. coli* BL21 (DE3) cells. The *E. coli* strains were grown overnight in the Luria–Bertani (LB) medium in the presence of 50 μg/mL ampicillin at 37°C. The overnight culture was diluted with fresh LB medium containing 50 μg/mL ampicillin and incubated at 37°C. Protein expression was induced at the growth point with an OD_600_ of 0.4–0.6 in the presence of 1 mM IPTG for 24 h at 15°C. Cells were harvested, washed twice with phosphate-buffered saline (PBS) (10 mM Na_2_HPO_4_, 2 mM NaH_2_PO_4_, 3 mM KCl, and 137 mM NaCl), and stored at 80°C until needed.

### Purification of SpeMreB

SpeMreB1, SpeMreB4, SpeMreB5, TmMreB, and their variants were purified by Ni^2+^-NTA affinity chromatography and gel filtration as previously described (27, 33, 49) independent of fusion with PrS. Briefly, cell pellets were resuspended in His trap buffer A (50 mM Tris-HCl pH 8.0, 300 mM NaCl, 50 mM imidazole-HCl pH 8.0), centrifuged at 12,000 ×*g* for 30 min at 4°C, and purified using HisTrap HP 5 mL (Cytiva; Tokyo, Japan). The imidazole concentration in the elution buffer for SpeMreB1 was changed from that used for SpeMreB5 (230 mM) to 500 mM because of the presence of double 6×His tags. The purified SpeMreBs were further subjected to HiLoad 26/600 Superdex 200 pg (Cytiva) in the gel filtration buffer (20 mM Tris-HCl pH 8.0, 300 mM NaCl) at 4°C. PrS was purified by Ni^2+^-NTA affinity chromatography using the same procedure as that used for SpeMreBs. Protein concentrations were determined from the absorbance at 280 nm measured using a NanoDrop One (Thermo Fisher Scientific), with the absorption coefficients estimated from ProtParam, as previously described (49).

### Electron microscopy

A 4 μL sample drop was placed onto a 400-mesh copper grid coated with carbon for 1 min at room temperature (23–27°C), washed with 10 μL water, stained with 2% (w/v) uranyl acetate for 45 s, air dried, and observed under a JEOL JEM-1010 transmission electron microscope (Tokyo, Japan) at 80 kV equipped with a FastScan-F214T charge-coupled device camera (TVIPS; Gauting, Germany). The image averaging of PrS-SpeMreB1 sheet was performed by CryoSPARC ver. 4.6.0. (50) with manually selected 1,284 particles. The major axis of PSMB1v was estimated by the distance measurement tool of ImageJ (National Institutes of Health; http://rsb.info.nih.gov/ij/).

### Sedimentation assays

Samples were prepared in the previously described buffer S (20 mM Tris-HCl pH 8.0, 1 M NaCl, 200 mM L-arginine-HCl pH 8.0, 5 mM DTT, 2 mM MgCl_2_ and 2 mM of the desired nucleotide), which was used for evaluating the polymerization activities of SpeMreB3 and SpeMreB5 (27). The reliability of the buffer S was confirmed by the sedimentation assay of TmMreB, where the band pattern was consistent with a previous sedimentation assay using KMEI buffer, which is commonly used for actin polymerization (Fig. S2H) (43). Co-sedimentation assays containing two of SpeMreB1, SpeMreB5, and PrS in the same sample solution were performed using this method. Sample incubation was initiated by mixing the sample solution with the desired protein concentration in gel filtration buffer and a reaction premix containing the other reagents in buffer S. After incubation at room temperature for 3 h, which was long enough to reach the equilibrium state of the polymerizations as confirmed by SpeMreB3 and SpeMreB5 (27), the samples were subjected to ultracentrifugation at 100,000 rpm at 23°C using a TLA-100 rotor (Beckman Coulter). The centrifugation time was set to 90 min (for samples containing PrS-SpeMreB1 variants) or 120 min (for samples without PrS-SpeMreB1), which is short enough to minimize the precipitation of SpeMreB monomers unless otherwise stated. After removal of the supernatant, the pellet was resuspended in 200 μL of water. The supernatant and pellet fractions were analyzed by SDS-PAGE on a 12.5% Laemmli gel and stained with Coomassie Brilliant Blue R-250. Of note, when preparing PrS-SpeMreB1-containing samples for SDS-PAGE, we omitted the steps of heat shock (at 95°C for 3 min in our laboratory protocol) or overnight incubation (at room temperature for more than 7 h in our laboratory protocol) after mixing with an SDS-PAGE sample solution (5% [v/v] glycerol, 0.025% [w/v] bromophenol blue, 62.5 mM Tris-HCl pH 6.8, 2.5% [w/v] sodium dodecyl sulfate, 5% [v/v] β-mercaptoethanol in our laboratory recipe), which are widely used for protein denaturation (51), because we have the following two experiences; (I) these steps stimulate the aggregation of PrS-SpeMreB1 in the SDS-PAGE sample solution and smearing of PrS-SpeMreB1 bands (Fig. S2I); and (II) all proteins used in this study were denatured immediately after mixing with the SDS-PAGE sample solution, as confirmed by the small amounts of smeared bands (see gel images in this paper, except Fig. S2I, in which the SDS-PAGE samples were prepared by the 95°C heat shock). The concentrations of the supernatant and pellet fractions were estimated as the product of the total SpeMreB concentration, and the ratio of each fraction to the sum of the supernatant and pellet fractions was measured using the ImageJ. The critical concentration was estimated as the x-intercept of the linear fit with the amount of precipitate at steady-state polymerization versus the total SpeMreB concentration.

### P_i_ release assays

P_i_ release measurements were performed as previously described in a standard buffer for SpeMreB polymerization (20 mM Tris-HCl pH 7.5, 100 mM KCl, 5 mM DTT, 2 mM MgCl_2_, and 2 mM ATP) (27, 49). Briefly, P_i_ release was detected using 2-amino-6-mercapto-7-methylpurine riboside, a molecular probe for P_i_ that reacts with P_i_ to produce a molecule that absorbs light at 360 nm.

### Circular dichroism (CD)

CD was measured for a 200 μL sample in a 1 mm thick quartz cuvette by J720W (JASCO; Tokyo, Japan) with a wavelength range of 200–250 nm under a nitrogen gas flow of 6 L/min at 20°C maintained by a temperature stabilizer. The measured CD intensities were converted to the mean residue ellipticities (*MRE*) using the following equation:

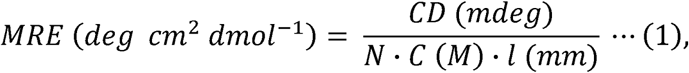

where *N* is the number of amino acid residues in the protein of interest, *C* is the protein concentration (*M*), and *l* is the path length of the cuvette used for measurements (mm). Secondary structure contents were estimated using BeStSeL (52–54) and compared with those estimated from the amino acid sequences using PSIPRED4 (55).

### Size-exclusion chromatography to study the interactions between SpeMreB1 and SpeMreB5

The samples were mixed in 20 mM Tris-HCl pH 7.5, 200 mM KCl and incubated on ice for 1 h. A 100 μL sample solution was loaded onto Superdex^TM^ 200 Increase 3.2/300 (Cytiva) equilibrated by 20 mM Tris-HCl pH 7.5, 200 mM KCl at 4°C. Protein signals were detected by measuring the absorbance at 280 nm.

### Protein expression in JCVI-syn3B

The construct for transferring *spemreB1* and *spemreB5* into the genome of *Mycoplasma mycoides* JCVI-syn3B (GenBank: CP069345.1) was assembled using pSD079 (56) and three DNA fragments: 496,781–498,064, including *spemreB1* and its promoter; 1,353,199–1,353,517, including the promoter of *spemreB4*; and 1,354,572–1,355,801, including *spemreB5* and its terminator. These fragments were amplified from the *S. eriocheiris* genome (GenBank: GCA_002028345.1) (57). Constructs for *spemreB1* E275R and *spemreB5* were prepared by inducing a mutation in pSD079 carrying *spemreB1* WT and *spemreB5* WT. pSD128 was used as a control strain carrying the puromycin resistance gene *puroR*. Each construct was transferred into JCVI-syn3B (58) (59) as previously described (15). These strains were cultured in the SP4 medium at 37°C to 0.03–0.04 at OD_620_. For protein profiling, the cell suspensions were concentrated 20-fold after washing with PBS containing sucrose (PBS/S) (75 mM sodium phosphate [pH 7.3], 68 mM NaCl, and 20 mM sucrose) and analyzed by SDS-PAGE. Protein bands were identified as previously described (38).

### Optical microscopy and video analyses of JCVI-syn3B cells

The cultures were centrifuged at 9000 ×*g* for 8 min at 10°C. The cell pellets were resuspended in PBS/S at one-fifth the volume of the culture, mixed with methylcellulose in PBS/S to a final concentration of 0.2%, and inserted into the tunnel slides. After 5 min at 25°C, 0.4% methylcellulose in PBS/S was inserted into the tunnel slide to wash out floating cells, and cells remaining near the glass surface were observed using an inverted microscope IX71 (Olympus; Tokyo, Japan) equipped with a UPlanSApo 100 × 1.4 numerical aperture (NA) Ph3 and complementary metal-oxide-semiconductor (CMOS) camera, DMK33UX174 (The Imaging Source Asia Co., Ltd. Taipei, Taiwan). The video was analyzed using the empirical gradient threshold (EGT) plugin (60) and a color footprinting macro (61) of ImageJ ver.1.53f51 (Fiji) (62).

### Liposome preparation

Chloroform solutions of 10 mg/mL DOPG (840475C, Avanti Polar Lipid, Alabaster, AL, USA), DOPC (850375C, Avanti Polar Lipid), SM (860062C, Avanti Polar Lipid), and CL (710335C, Avanti Polar Lipid) were used for liposome construction. For SpiroLipid liposomes, chloroform solutions of these lipids were mixed in a previously described molar ratio (17, 63) (DOPG:DOPC:SM:CL = 0.38:0.14:0.33:0.15). For liposome preparation, a chloroform solution with the desired lipid composition was aliquoted into a clean glass bottle and was completely dried by N_2_ gas flow and a vacuum desiccation at room temperature. The remaining lipid layer was hydrated at room temperature using a standard buffer with increasing KCl concentration to 200 mM (to reduce the background precipitation of SpeMreBs) containing 1 mM MgCl_2_ (for efficient liposome construction) and subjected to water bath sonication for 1 min. The liposome solution was extruded at room temperature using a 100 nm polycarbonate membrane (610005, Avanti Polar Lipid). The liposome solution was stored at 4°C and used within 2 days from the construction.

### Liposome binding assays

SpeMreB samples were dialyzed to 20 mM Tris-HCl pH 7.5, 200 mM KCl, 1 mM MgCl_2_ at 4°C for overnight. The dialyzed samples were centrifuged at 20,000 ×*g* for 10 min at 4°C to remove aggregates, mixed with a liposome solution with a final SpeMreB concentration and sample volume of 2 μM and 200 μL, respectively, and incubated for 15 min at room temperature. For liposome binding assays in the presence of a nucleotide, SpeMreBs were incubated in the presence of a nucleotide for 5 min at room temperature before liposome addition at twice the SpeMreB and nucleotide concentrations (4 μM and 2 mM, respectively). The incubated samples were centrifuged at 48,000 rpm (approximately 100,000 ×*g*) for 30 min at 23°C using a TLA-100 rotor. The supernatant was removed, and the pellet was resuspended in water. SpeMreB concentrations in each fraction were estimated by SDS-PAGE and subsequent image analysis as described in “Sedimentation assays” in the Materials and Methods section.

## Acknowledgments

We thank Ms. Junko Shiomi, Ms. Tomomi Shimonaka, Mr. Yuhei O Tahara, Dr. Ritsuko Fujii, and Dr. Yoshihiro Yamaguchi (all of whom are affiliated with the Graduate School of Science, Osaka Metropolitan University, Japan) for their technical assistance with SpeMreB construction and expression, technical assistance with MALDI-TOF MASS spectrometry, technical assistance for negative-staining EM, providing a facility for CD measurements, and pCold-PrS. We thank Prof. Robert C. Robinson (Research Institute for Interdisciplinary Science, Okayama University, Okayama, Japan and School of Biological Science and Engineering, Vidyasirimedhi Institute of Science and Technology, Thailand) and Dr. Yoshihito Kitaoku (Research Institute for Interdisciplinary Science, Okayama University, Okayama, Japan) for providing the construct for expressing TmMreB and developing the environment for using CryoSPARC, respectively.

## Author contributions

Conceptualization: DT and MM. Methodology: DT, HK, and HM. Investigation: DT and HK. Visualization: DT and HK. Supervision: IF and MM. Writing the original draft: DT, MM, and IF. Writing—review and editing: DT, HK, HM, MM, and IF. Funding acquisition: DT, HK, and MM.

## Data and materials availability

Raw data are available from the corresponding author upon reasonable request.

---

**Table S1.** Results of peptide mass fingerprinting of bands shown in Fig. 4C. Roman numbers indicate protein bands shown in Fig. 4C. All proteins were detected in both peptide mass fingerprinting and MS/MS analyses. The Mascot protein score is defined by −10×log(P), where P is the probability that the observed match is a random event. The reference dataset for peptide mass finger printing was prepared by adding the amino acid sequence of SpeMreB1 E275R to the coding sequence libraries of Synthetic bacterium JCVI-syn3B (taxonomy ID: 2806337) and *Spiroplasma eriocheiris* (taxonomy ID: 743698) in the NCBI database. The protein with star was identified in the MS/MS analysis, although its score in the peptide mass finger printing

| Protein band | Protein ID | Gene name | Annotation | Mascot protein score | Mass [kDa] | Sequence coverage [%] | Species |
| --- | --- | --- | --- | --- | --- | --- | --- |
| i | QWN46562.1 | *gap* | Glyceraldehyde-3-phosphate dehydrogenase | 39 | 37.2 | 24 | Synthetic bacterium JCVI-Syn3B |
| ii | QWN46586.1 | *rpoA* | DNA-directed RNA polymerase subunit alpha | 60 | 35.0 | 45 | Synthetic bacterium JCVI-Syn3B |
| iii | QWN46506.1* | *ldh* | L-lactate dehydrogenase | 20 | 34.8 | 26 | Synthetic bacterium JCVI-Syn3B |
| iv | AHF58343.1 | *mreB5* | Cell shape-determining protein MreB | 69 | 38.7 | 28 | *Spiroplasma eriocheiris* |
| v | AHF57598.1 | *mreB1* | Cell shape-determining protein MreB | 53 | 38.0 | 34 | *Spiroplasma eriocheiris* |
| vi | AHF58343.1 | *mreB5* | Cell shape-determining protein MreB | 51 | 38.7 | 26 | *Spiroplasma eriocheiris* |
| vii | AHF57598.1-2 | *mreB1 E275R* | Cell shape-determining protein MreB | 59 | 38.0 | 30 | *Spiroplasma eriocheiris* |

\* Although this protein did not get the highest score in the peptide mass finger printing, it was identified in the MS/MS analysis.

**Fig. S1.**
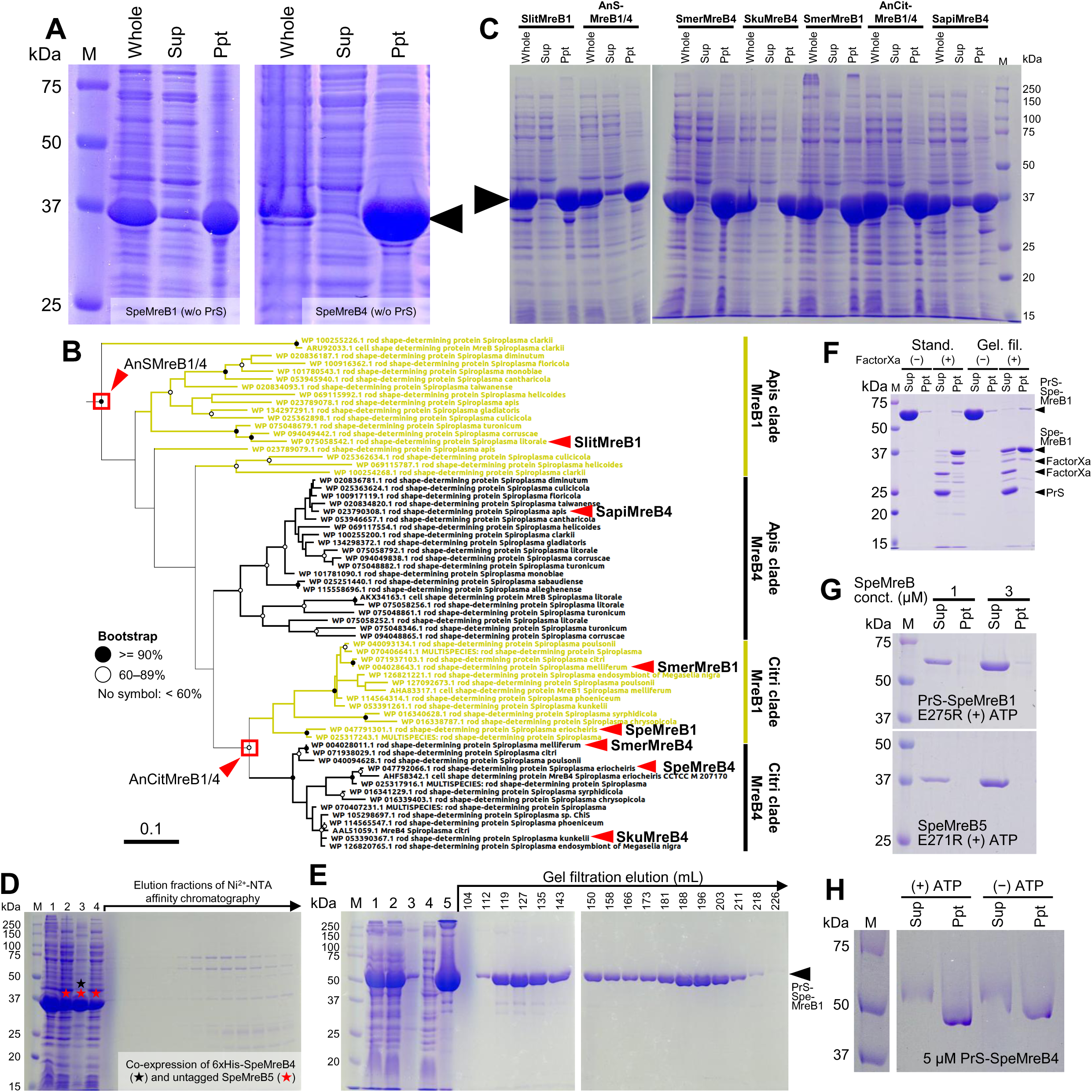
Preparation and evaluation of PrS-SpeMreB1 and PrS-SpeMreB4. For gel images, protein size standards were visualized in lane M with the molecular masses next to each band. (**A**) Expression of SpeMreB1 WT (left) and SpeMreB4 WT (right) without the PrS tag. The SpeMreBs were expressed by *E. coli* BL21 (DE3) as a 6×His-tag fusion at each N-terminus using the pCold-15b expression system. Cells were sonicated in 50 mM Tris-HCl pH 8.0, 300 mM NaCl, and centrifuged at 12,000 ×*g* for 30 min at 4°C. The whole cell lysate, supernatant, and pellet fractions were analyzed by SDS-PAGE. The band positions for SpeMreB1 WT and SpeMreB4 WT are indicated by an arrowhead. (**B**) Phylogenetic tree of SMreB1 and SMreB4. The scale bar is in units of the number of amino acid substitutions per site. Sequences used for expression experiments in this study are indicated by red triangles with their labeling. (**C**) Expression of *S. melliferum* MreB4 (SmerMreB4), *S. kunkelii* MreB4 (SkuMreB4), *S. melliferum* MreB1 (SmerMreB1), *S. apis* MreB4 (SapiMreB4), *S. litorale* MreB1 (SlitMreB1), a reconstituted ancestor of the Citri clade MreB1 and MreB4 (AnCitMreB1/4), and a reconstituted ancestor of *Spiroplasma* MreB1 and MreB4 (AnSMreB1/4). These constructs were cloned, expressed, isolated, and analyzed by SDS-PAGE using the same procedures as those in Fig. S1A. The band positions of these constructs are indicated by an arrowhead. (**D**) Co-expression of 6×His-tagged SpeMreB4 WT and untagged SpeMreB5 WT and their purification. *E. coli* cells co-expressing these SpeMreB4 and SpeMreB5 constructs were sonicated in 50 mM Tris-HCl pH 8.0, 50 mM Imidazole-HCl pH 8.0, 300 mM NaCl, centrifuged at 12,000 ×*g* for 30 min at 4°C and applied to a Ni^2+^-NTA affinity chromatography column. The following fractions were visualized by a Coomassie-stained 12.5% Laemmli gel; fraction 1: lysate of *E. coli* co-expressing SpeMreB4 WT and SpeMreB5 WT; fractions 2 and 3: soluble (2) and insoluble (3) fractions of the whole cell lysate (1); fraction 4: flow-through fraction of Ni^2+^-NTA affinity chromatography; other fractions: elution fractions of Ni^2+^-NTA affinity chromatography with a gradually increasing imidazole concentration. Red and black stars indicate the bands in which SpeMreB4 and SpeMreB5 were detected, respectively, by the peptide mass fingerprinting technique using MALDI-TOF MASS spectrometry. SpeMreB5 was detected in lanes 2, 3, and 4, whereas SpeMreB4 was detected in lane 3 and not detected in lane 2, indicating that SpeMreB4 remained insoluble despite co-expression with SpeMreB5. (**E**) Purification procedure of PrS-SpeMreB1 WT. The following fractions were visualized using a Coomassie-stained 12.5% Laemmli gel; fraction 1: lysate of *E. coli* expressing PrS-SpeMreB1 WT; fractions 2 and 3: soluble (2) and insoluble (3) fractions of the whole cell lysate (1); fractions 4 and 5: flow-through (4) and elution (5) fractions of Ni^2+^-NTA affinity chromatography; the other fractions: gel filtration elution fractions with the elution volume in Fig. 1B indicated on each lane. The band position of PrS-SpeMreB1 is indicated by an arrowhead. (**F**) Removal of the PrS tag from PrS-SpeMreB1 WT. PrS-SpeMreB1 WT at 10 μM concentration was dialyzed against 20 mM Tris-HCl pH 7.5, 100 mM KCl (Stand.) or 20 mM Tris-HCl pH 8.0, 300 mM NaCl (Gel. fil.) for 20 h at 4°C in the presence (+) or absence (−) of 0.1 mg / ml factorXa (Novagen, Merck Co. Ltd.) and centrifuged at 20,000 ×*g* for 10 min at 4°C. The resulting supernatant and pellet fractions were visualized by SDS-PAGE. The band positions of PrS-SpeMreB1 WT, SpeMreB1 WT, factorXa, and PrS are indicated by closed triangles. The other bands are probably due to non-specific cleavage of PrS-SpeMreB1 by factorXa. (**G**) Sedimentation assays of PrS-SpeMreB1 E275R (top) and SpeMreB5 E271R (bottom) in the presence of 2 mM ATP. (**H**) Sedimentation assays of 5 μM SpeMreB4 WT in the presence or absence of 2 mM ATP.

**Fig. S2.**
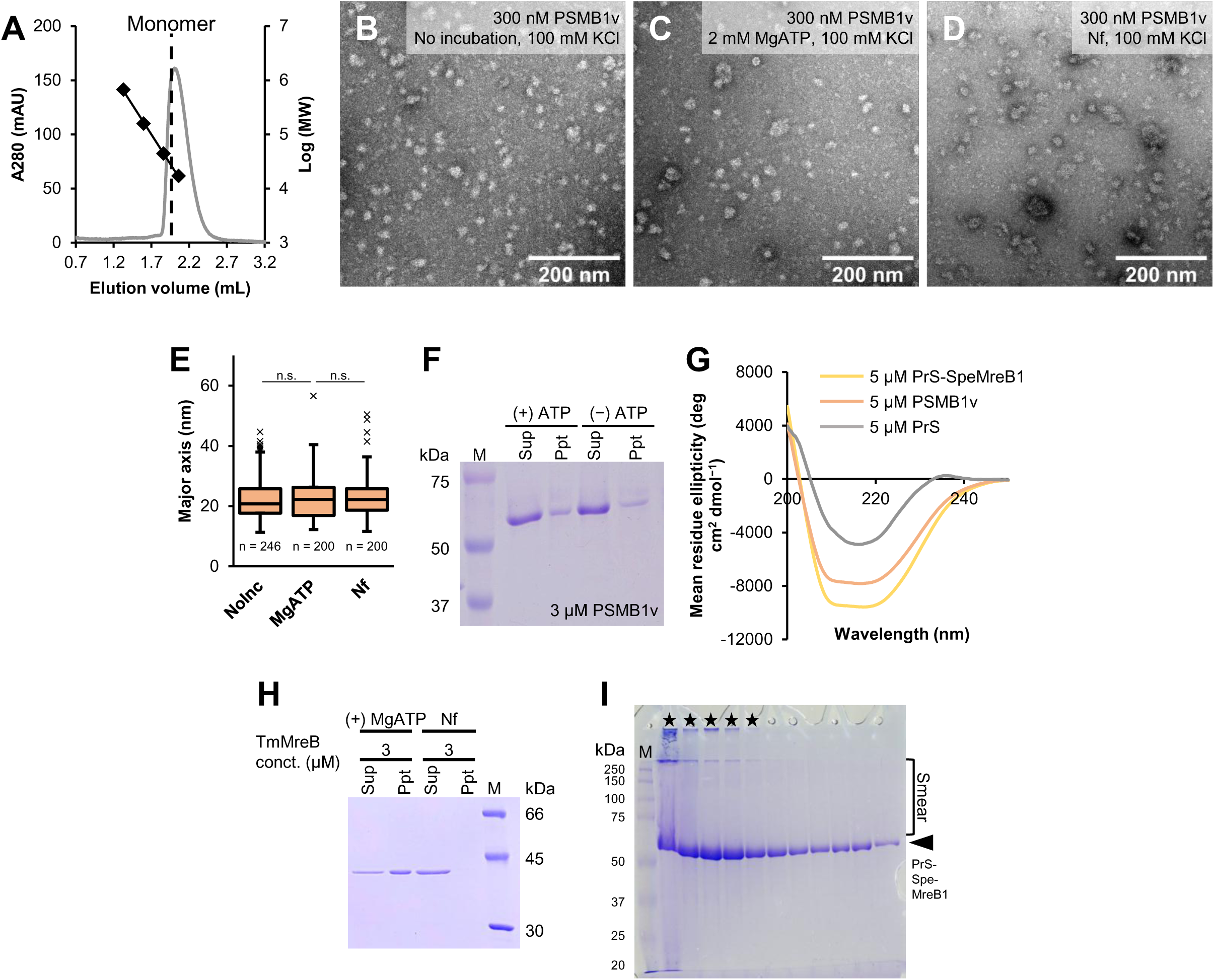
Evaluation of PSMB1v. For gel images, protein size standards were visualized in lane M with the molecular masses next to each band. (**A**) Gel filtration profiles of 50 μM PrS by Superdex^TM^ 200 Increase 3.2/300 in the gel filtration buffer. Bovine thyroglobulin (670 kDa), bovine γ-globulin (158 kDa), chicken ovalbumin (44 kDa), and horse myoglobin (17 kDa) were used as the protein size standards, and their elution volumes are plotted using closed diamonds with the linear fit over the log of their molecular weights. The estimated elution volume of the monomeric PrS (23 kDa) shown as a dashed line is in good agreement with the peak top position of the PrS elution pattern, indicating that most of PrS molecules were monomers at this concentration. (**B–D**) Negative staining EM images of 300 nM PSMB1v in the standard buffer (**B**) without an incubation and (**C–D**) with an incubation at RT for 1 h in the (**C**) presence and (**D**) absence of 2 mM MgATP. (**E**) The size distribution of PSMB1v at the conditions shown in Fig. S2B–D. The major axes of the all-detectable particles were measured for each micrograph and accumulated until approximately 200 data were collected. Sample numbers of each condition are indicated below each box. Values over the upper fence (75th percentile + 1.5 × (75th percentile – 25th percentile)) are defined as outliers and plotted as black crosses. Symbols indicate *p*-value supported by Student’s *t-*test (n.s. *p* > 0.05). (**F**) Sedimentation assays of 3 μM PSMB1v WT in the presence or absence of 2 mM ATP. (**G**) CD spectra of 5 μM PrS-SpeMreB1 WT (yellow), 5 μM PSMB1v WT (orange), and 5 μM PrS (gray) in the gel filtration buffer. (**H**) Sedimentation assays of 3 μM TmMreB in the presence or absence of 2 mM ATP. (**I**) An example of an SDS-PAGE profile of PrS-SpeMreB1 WT in which the samples were prepared with a heat shock step. After mixing with the SDS-PAGE sample solution, PrS-SpeMreB1 in the gel filtration buffer was heated at 95°C for 3 min before loading onto the SDS-PAGE gel, and smears of PrS-SpeMreB1 bands appeared at the region indicated by half brackets. Black stars indicate the lanes in which multiple amounts of PrS-SpeMreB1 WT were stacked on top of stacking and running gels. The band position for unaggregated PrS-SpeMreB1 WT is indicated by an arrowhead.

**Fig. S3.**
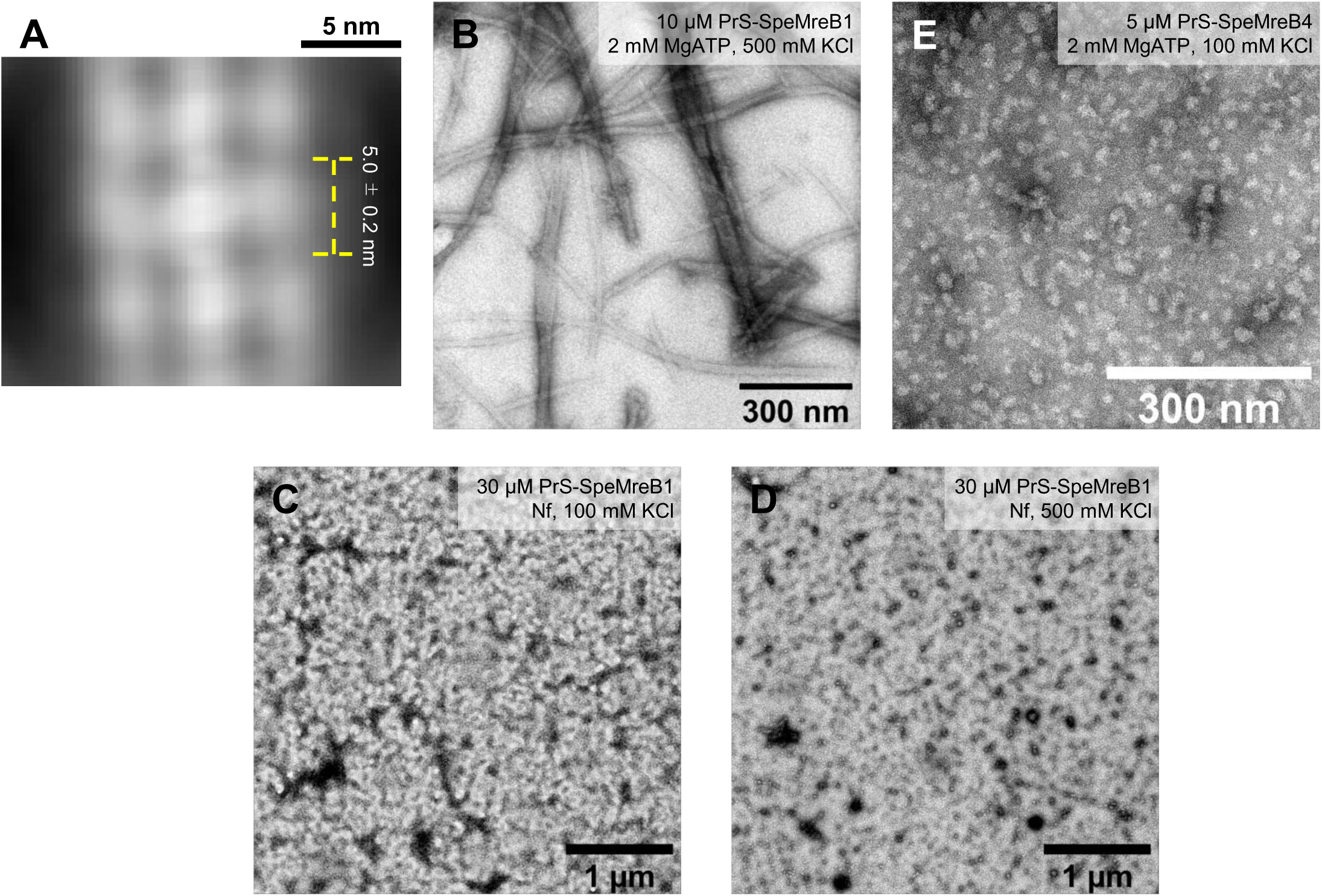
Negative staining EM images of PrS-SpeMreB1 and PrS-SpeMreB4. (**A**) Two-dimensionally averaged image of PrS-SpeMreB1 sheet at the condition of Fig. 1C from 881 particles. The subunit repeat is estimated to be 5.0 ± 0.2 nm. (**B–E**) Negative staining EM images of (**B**) 10 μM and (**C–D**) 30 μM PrS-SpeMreB1 WT and (**E**) 5 μM SpeMreB4 WT in (**C** and **E**) the standard buffer and (**B** and **D**) that with the increasing KCl concentration of 500 mM in the (**B** and **E**) presence or (**C–D**) absence of 2 mM MgATP.

**Fig. S4.**
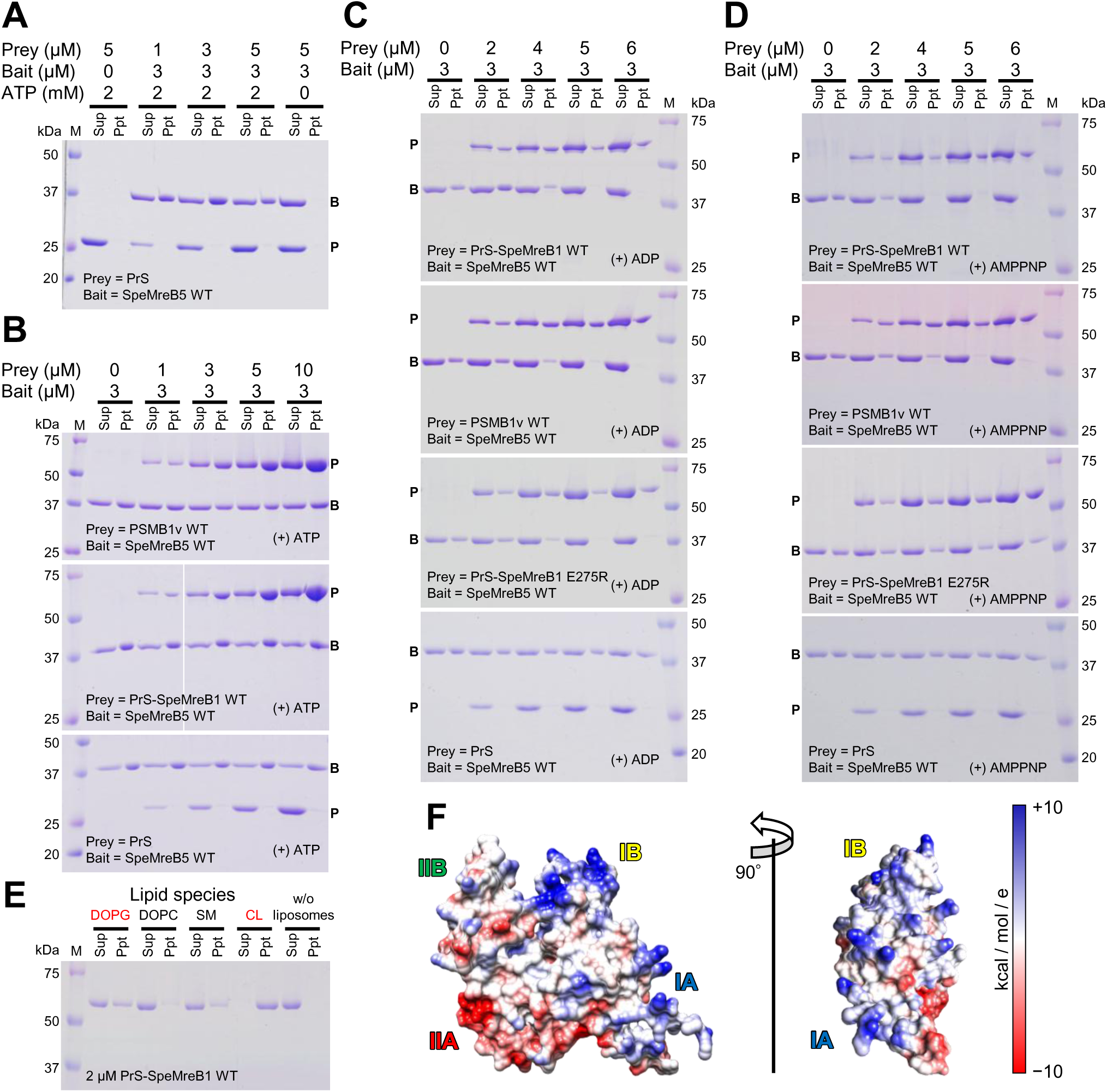
Sedimentation assays of PrS-SpeMreB1 variants. For gel images, protein size standards were visualized in Lane M with the molecular masses beside each band. The band positions of prey and bait in gel images of co-sedimentation assays are indicated by P and B, respectively. (**A**) Co-sedimentation assays of 3 μM SpeMreB5 WT and 1 μM to 5 μM PrS in the presence or absence of 2 mM ATP. (**B**) Co-sedimentation assays of 3 μM SpeMreB5 WT and 0 μM to 10 μM PSMB1v WT (top), PrS-SpeMreB1 WT (middle), or PrS (bottom) in the presence of 2 mM ATP. Pellet amounts of SpeMreB5 were estimated and summarized in Fig. 3B. (**C–D**) Co-sedimentation assays of 3 μM SpeMreB5 WT and 0 μM to 6 μM PrS-SpeMreB1 WT (top), PSMB1v WT (second top), PrS-SpeMreB1 E275R (second bottom), or PrS (bottom) in the presence of 2 mM (**C**) ADP or (**D**) AMPPNP. Pellet amounts of SpeMreB5 were estimated and summarized in Fig. 3B. (**E**) Liposome binding assays of 2 μM PrS-SpeMreB1 WT in the presence of 1 mM liposomes composed of 100% DOPG, DOPC, SM, or CL. Negatively charged lipids (DOPG and CL) are indicated by red characters. (**F**) The surface representation of the full-length SpeMreB1 monomer modeled by AlphaFold2 (41). The Coulombic electrostatic potential is indicated by the color gradient from blue (+10 kcal/mol/e) to red (−10 kcal/mol/e) via white (0 kcal/mol/e). Subdomains IA, IB, IIA, and IIB are indicated at the corresponding positions of the structure. The side view of the membrane binding region on most MreBs (21, 28) is shown on the right beside the front view on the left.

**Movie S1. Combined real-time movies (10 s) of three syn3B strains collected from phase contrast.** The strain names are shown at the top left of each panel. The scale bar indicates the bottom of the right panel.

**Supplementary data 1. Nucleotide sequence of multiple cloning sites in pCold-PrS.** Sequences of PrS, 6xHis-tag, factor Xa cleavage site, NdeI site, and BamHI site are shown in gray, blue, brown, purple, and red, respectively. The 5’- and 3’-ends are respectively connected with 5’- and 3’-UTR of a cold shock gene, *cspA*. ATGAATCATAAAGTG**CATCATCATCATCATCACATGGCAAATATTACCGTTTTCTA TAACGAAGACTTCCAGGGTAAGCAGGTCGATCTGCCGCCTGGCAACTATACCCGCGCCCAGTTGGCGGCGCTGGGCATCGAGAATAATACCATCAGCTCGGTGAAGGTGCCGCCTGGCGTGAAGGCTATCCTGTACCAGAACGATGGTTTCGCCGGCGACCAGATCGAAGTGGTGGCCAATGCCGAGGAGTTGGGCCCGCTGAATAATAACGTCTCCAGCATCCGCGTCATCTCCGTGCCCGTGCAGCCGCGCATGGCAAATATTACCGTTTTCTATAACGAAGACTTCCAGGGTAAGCAGGTCGATCTGCCGCCTGGCAACTATACCCGCGCCCAGTTGGCGGCGCTGGGCATCGAGAATAATACCATCAGCTCGGTGAAGGTGCCGCCTGGCGTGAAGGCTATCCTCTACCAGAACGATGGTTTCGCCGGCGACCAGATCGAAGTGGTGGCCAATGCCGAGGAGCTGGGTCCGCTGAATAA TAACGTCTCCAGCATCCGCGTCATCTCCGTGCCGGTGCAGCCGAGG**GGTACC**AT TGAAGGCCGCCATATG**GTCGACCTCGAG**GGATCC**CGTGGTGAAATC**CATCACCAT CACCATCAC**TAATCTAGATAGGTAA

**Supplementary data 2. Amino acid sequences and accession IDs of SMreB1 and SMreB4 used in expression experiments (Fig. S1B–C).** Accession IDs are summarized for sequences deposited in GenBank. The amino acid sequences are summarized for the other sequences.

>SpeMreB1: WP_047791301.1

>SpeMreB4: WP_047792066.1

>SmerMreB1: WP_004028643.1

>SmerMreB4: WP_004028011.1

>SkuMreB4: WP_053390367.1

>SapiMreB4: WP_023790308.1

>SlitMreB1: WP_075058542.1

>AnCitMreB1/4

MALFNSKKPTFVSMDLGTANTLVYVSGSGIVYNEPSIVAYKIKENRIIAVGNEAYKMIGKGNKSIRIVRPMVDGVITDIRATEAQLRYIFNKLRISKQLKNSIMLLACPSVITELEKNALKKIAMNLGADKVFVEEEVKMAALGGGVDIYKPAGNLVVDMGGGTTDIAVLASGDIVLSKSVKVAGNYLNDEIQKFIRSQYGLEIGIKTAEQIKIEIGSLAKYPDERKMKVYGRDVVSGLPREIEVTPEEIREVLKVPVSRIIDLTVQVLEETPPELAGDIFRNGITICGGGALIKGIDKYFEDTLQLPAKIGEQPLLAVINGTKKFESDIYDILRQEHMHTKELNY

>AnSMreB1/4

MAKKKSKKPTFVSMDLGTANTLVYISGQGIVYNEPSIVAYKIKENKIIAVGEEAYKMIGKGNKNIRIVRPMVDGVITDIRATEAQLRYIFNKLRISKTLKNSIMLLACPSVITELEKNALKKIAMNLGADKVFVEEEVKMAALGGGVNIYAPTGNLVVDMGGGTTDIAVLASGDIVLSKSIKVAGNYLNDEIQKFIRSQYGLEIGIKTAEQIKINIGSLAKYPDERKMKVYGRDVVSGLPREIEITPEEIREVLKVPVSRIIDLTVQVLEETPPELAGDIFRNGITICGGGALIKGIDKYFEDTLQ LPTKIGEQPLLAVINGTKKFESDIYDILKEEHNHTKELNY

